# Explicit expression of mesophyll conductance in the traditional leaf photosynthesis–transpiration coupled model and its physiological significances

**DOI:** 10.1101/2021.09.22.461327

**Authors:** Hong Luo, Marc Carriquí, Miquel Nadal, Tuo Han, Christiane Werner, Jian-feng Huang, Jiao-lin Zhang, Zhi-guo Yu, Feng-min Li, Xiang-wen Fang, Wei Xue

## Abstract

- Almost all terrestrial biosphere models (TBMs) still assume infinite mesophyll conductance (*g*_m_) to estimate photosynthesis and transpiration. This assumption has caused low accuracy of TBMs to predict leaf gas exchange under certain conditions.
- In this study, we developed a photosynthesis-transpiration coupled model that explicitly considers *g*_m_ and designed an optimized parameterization solution through evaluating four different *g*_m_ estimation methods in 19 C_3_ species at 31 experimental treatments.
- Results indicated that temperature responses of the maximum carboxylation rate (*F*_cmax_) and the electron transport rate (*J*_max_) estimated by fusing the Bayesian retrieval algorithm and the Sharkey online calculator together with *g*_m_ temperature response estimated by fusing the chlorophyll fluorescence-gas exchange method and anatomy method predicted leaf gas exchange more accurately. The *g*_m_ temperature response exhibited activation energy (Δ*H*_a_) of 63.13 ± 36.89 kJ mol^-1^ and entropy (Δ*S*) of 654.49 ± 11.36 J K^-1^ mol^-1^. The *g*_m_ optimal temperature (*T*_opt__g_m_) explained 58% of variations in photosynthesis optimal temperature (*T*_optA_). The *g*_m_ explicit expression has equally important effects on photosynthesis and transpiration estimations.
- Results advanced understandings of better representation of plant photosynthesis and transpiration in TBMs.

## Introduction

The leaf photosynthesis-transpiration coupled model is the basis of vegetation photosynthesis and transpiration estimations performed using terrestrial biosphere models (TBMs) (Dai *et al*., 2003; Rogers *et al*., 2017). The model is based on the biochemical-based photosynthesis theory developed by Farquhar *et al.* (1980), which considers the leaf stomatal aperture as the primary gauge of CO_2_ influx and water vapor efflux and assumes an infinite conductance of CO_2_ diffusion from sub-stomatal cavities to the chloroplast stroma (i.e. mesophyll conductance, *g*_m_). In such cases, there is no CO_2_ concentration drawdown between the intercellular airspace and the chloroplast stroma. However, now it is widely accepted not only that there is a significant CO_2_ diffusion drawdown from the intercellular airspace to the chloroplast stroma, but that the resistance that causes this drawdown can be similar or greater than stomatal resistance in land plants (Flexas *et al*., 2012; Evans and von Caemmerer, 2013; von Caemmerer and Evans, 2015; Gago *et al*., 2019). Currently, almost all TBMs do not explicitly consider *g*_m_ (Rogers *et al*., 2017; Knauer *et al*., 2019; 2020; Iqbal *et al*., 2021), primarily because of (1) huge disputes in response characteristics of *g*_m_ to ambient environment, particularly the temperature response (von Caemmerer and Evans, 2015; Bahar *et al*., 2018; Shrestha *et al*., 2019; Li *et al*., 2020; Evans, 2021), and (2) lack of a *g*_m_ finite photosynthesis-transpiration coupled model that can be applied to as many C_3_ species as possible (Niinemets *et al*., 2009; Xue *et al*., 2017; Knauer *et al*., 2020).

Mesophyll conductance cannot be directly measured mainly because it is not possible to determine the CO_2_ concentration within the chloroplast stroma. Thus, modeling methods are required to estimate *g*_m_, which could be grouped into four classes according to the algorithm principle and field measurement tools: the chlorophyll fluorescence-gas exchange method (i.e. *g*_m_F_) (Harley *et al*., 1992); the anatomy method (i.e. *g*_m_A_) (Tosens *et al*., 2012; Tomas *et al*., 2013); the ^13^C isotope discrimination method (Evans *et al*., 1986; Barbour *et al*., 2007; Pons *et al*., 2009; Evans and von Caemmerer, 2013; Flexas *et al*., 2013); and the curve-fitting parameter retrieval methods (Sharkey *et al*., 2007; von Caemmerer *et al*., 2009; Yin and Struik, 2009; Gu *et al*., 2010; Zhu *et al*., 2011; von Caemmerer, 2013, Han *et al*., 2020; Xiao *et al*., 2021). These methods have their own advantages and weakness in terms of model parameter assumptions made during estimation, resulting in large discrepancies in the estimated temperature response of *g*_m_ (Pons *et al*., 2009; Evans, 2021). Therefore, we decided to select the best *g*_m_ estimation method in terms of accuracy in predicting leaf gas exchange at any temperature, namely a good criterion to select at each temperature which *g*_m_ estimation method works best is the one that could predict leaf gas exchange more accurately.

Effects of the *g*_m_ finite expression on photosynthesis have been widely demonstrated, whereas few attentions are paid on transpiration. Considering the indirect effects of *g*_m_ on transpiration and the direct effect on photosynthesis through its control on chloroplast CO_2_ concentration, Knauer *et al*. (2020) determined that *g*_m_ has a greater effect on photosynthesis but no significant effects on transpiration in most species and hence proposed the ‘the asymmetric effects on photosynthesis and transpiration estimations’ hypothesis. Kumarathunge *et al.* (2019) found that variations in photosynthesis optimal temperature (*T*_optA_) can be primarily explained by changes in the ratio of the apparent maximum electron transport rate and the apparent maximum carboxylation rate at 25°C (*J*_a,25_/*V*_a,25_, *JV*r), thus proposing ‘the *JV*r biochemical limitations’ hypothesis. *J*_a,25_ and *V*_a,25_ estimations in their study were derived from the *A*_n_/C_*i*_ curve without explicitly considering *g*_m_. Previous results indicated significant changes in *T*_optA_ under different intercellular CO_2_ concentrations (*C*_i_) in C_3_ species (Farquhar *et al*., 1980; Rogers *et al*., 2017), suggesting that the CO_2_ substrate levels significantly affect *T*_optA_. The CO_2_ substrate levels inside the chloroplasts in turn are strongly controlled by *g*_m_. We speculated that the plausibility of the *T*_optA_-*JV*r relationship reported by Kumarathunge *et al.* (2019) is questionable in reality. The observed variations in *T*_optA_ among plant species may not be solely explained by biochemical limitations.

As a first step towards the better representation of plant photosynthesis and transpiration in most TBMs, we attempted to develop a *g*_m_ finite model that could be directly implemented in most TBMs, in addition to design an optimized parameter configuration solution that is physiologically meaningful for as many C_3_ species as possible. The optimized parameterization solution was designed by comparing multiple different parameter estimation methods used for *g*_m_, *V*_cmax_, and *J*_max_ estimations. The response features of *g*_m_, *V*_cmax_, and *J*_max_ to temperature were therefore determined. Dynamic changes in apparent *g*_m_ in response to irradiance were not of main consideration in the current study, since the underlying conductance at the cellular level may remain unchanged with varying light environments (Evans, 2021). Validation of the predictability of the *g*_m_ finite model compared to a traditional photosynthesis model assuming infinite *g*_m_ (abbreviated as the *g*_m_ infinite model) was performed in 19 C_3_ species under 31 experimental conditions. The effects of the *g*_m_ finite expression on leaf photosynthesis and transpiration estimations were quantified to evaluate the ‘the *JV*r biochemical limitations’ hypothesis proposed by Kumarathunge *et al.* (2019) and ‘the asymmetric effects on photosynthesis and transpiration estimations’ hypothesis proposed by Knauer *et al*. (2020).

## Materials and methods

### Model description

In line with the photosynthesis-transpiration coupled model adopted by most TBMs, the Farquhar, von Caemmerer & Berry (1980) (FvCB) photosynthesis model and a stomatal conductance sub-model (Ball *et al*., 1987; Leuning, 1995) were coupled to quantify leaf carbon uptake and water loss through transpiration. *A*_n_ is limited either by the Rubisco carboxylation rate at a low CO_2_ concentration (*W*_c_) or the RuBP regeneration rate at a relatively high CO_2_ concentration because of low electron transport rates (*W*_j_) or deficit inorganic phosphate for photophosphorylation (*W*_p_). *A*_n_ can be expressed as follows:

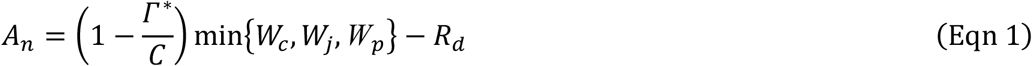

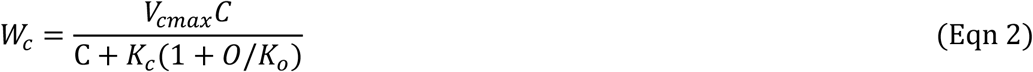

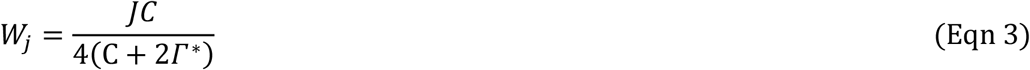

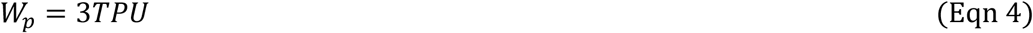

where *V*_cmax_ is the maximum carboxylation rate; *Γ*\* is the CO_2_ compensation point without mitochondrial respiration; *K*_c_ and *K*_o_ are the Michaelis-Menten constants for CO_2_ and O_2_, respectively; *R*_d_ and *J* are dark respiration in the light and the electron transport rate, respectively; *TPU* is the rate of triose phosphate export from the chloroplasts. *C* is *C*_i_ for the *g*_m_ infinite model or CO_2_ concentration inside the chloroplasts (*C*_c_) for the *g*_m_ finite models, which can be determined using Eqns. 9 and 10. *J* is modeled as a function of incident photosynthetically active radiation (*PAR*), which is calculated using the non-rectangle hyperbola equation (Eqn. 5) (Farquhar *et al*., 1980; Medlyn *et al*., 2002; Sharkey *et al*., 2007; Rogers *et al*., 2017; Kumarathunge *et al*., 2019) as follows:

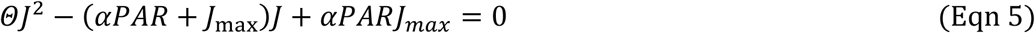

where *α* is the quantum yield of electron transport; *Θ* is the curvature of the non-rectangle hyperbola equation (Table 1).

**Table 1.**
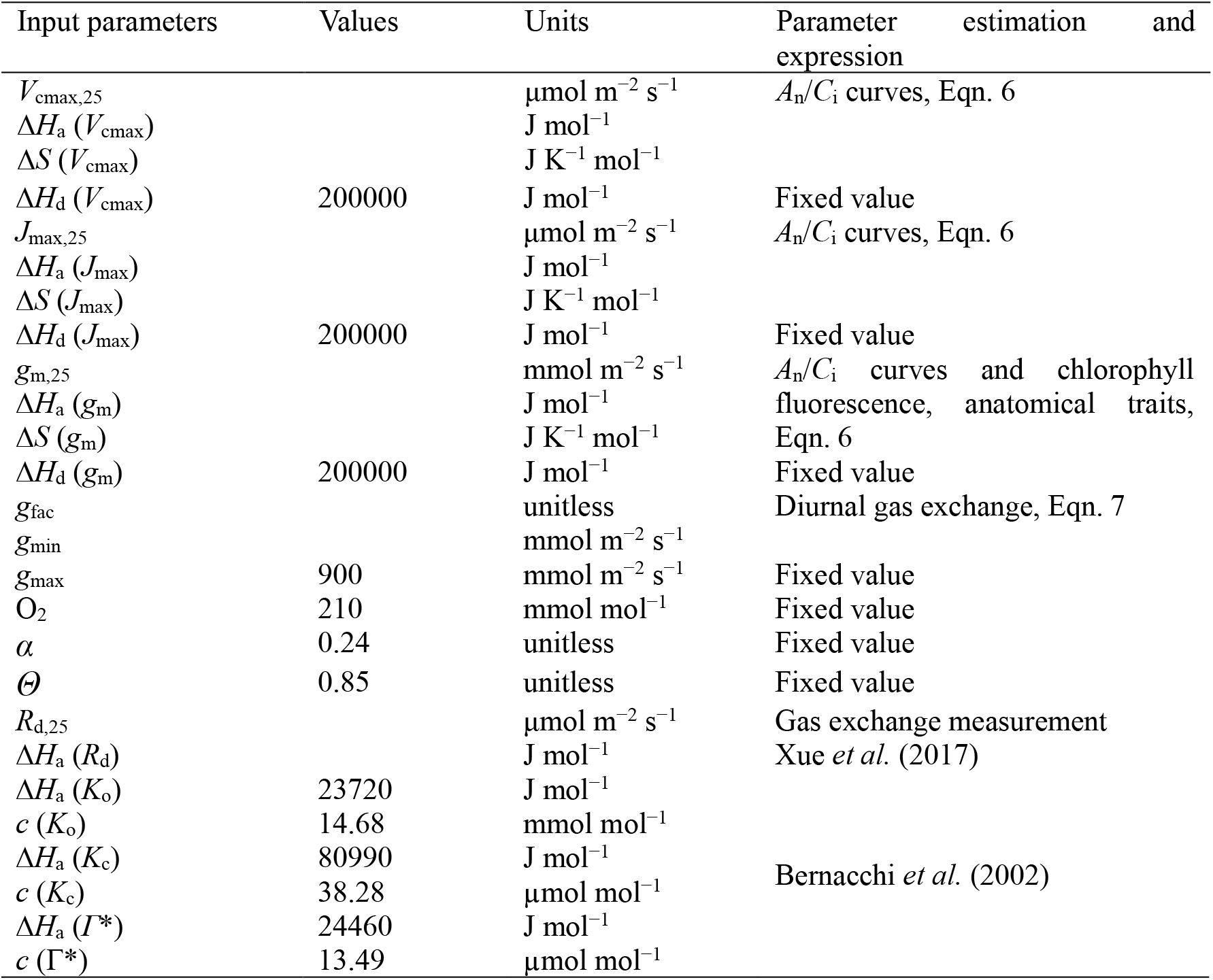
Input parameters of the *g*_m_ finite photosynthesis-transpiration coupled model. Fixed values were referred to Harley and Tenhunen (1991), Xue *et al.* (2017), Knauer *et al.* (2019), and Kumarathunge *et al.* (2019). The Rubisco kinetic parameters (*K*_c_, *K*_o_, and *Γ*\*) were modeled by the Arrhenius function, i.e. Γexp(*c*-Δ*H*_a_/*R*/*T*_k_)].

| Input parameters | Values | Units | Parameter estimation and expression |
| --- | --- | --- | --- |
| $V_{cmax,25}$ | | $\mu\text{mol m}^{-2} \text{ s}^{-1}$ | $A_n/C_i$ curves, Eqn. 6 |
| $\Delta H_a (V_{cmax})$ | | $\text{J mol}^{-1}$ | |
| $\Delta S (V_{cmax})$ | | $\text{J K}^{-1} \text{ mol}^{-1}$ | |
| $\Delta H_d (V_{cmax})$ | 200000 | J mol<sup>-1</sup> | Fixed value |
| $J_{max,25}$ | | μmol m<sup>-2</sup> s<sup>-1</sup> | $A_n/C_i$ curves, Eqn. 6 |
| $\Delta H_a (J_{max})$ | | J mol<sup>-1</sup> | |
| $\Delta S (J_{max})$ | | J K<sup>-1</sup> mol<sup>-1</sup> | |
| $\Delta H_d (J_{max})$ | 200000 | J mol<sup>-1</sup> | Fixed value |
| $g_{m,25}$ | | mmol m<sup>-2</sup> s<sup>-1</sup> | $A_n/C_i$ curves and chlorophyll |
| $\Delta H_a (g_m)$ | | J mol<sup>-1</sup> | fluorescence, anatomical traits, |
| $\Delta S (g_m)$ | | J K<sup>-1</sup> mol<sup>-1</sup> | Eqn. 6 |
| $\Delta H_d (g_m)$ | 200000 | J mol<sup>-1</sup> | Fixed value |
| $g_{fac}$ | | unitless | Diurnal gas exchange, Eqn. 7 |
| $g_{min}$ | | mmol m<sup>-2</sup> s<sup>-1</sup> | |
| $g_{max}$ | 900 | mmol m<sup>-2</sup> s<sup>-1</sup> | Fixed value |
| $O_2$ | 210 | mmol mol<sup>-1</sup> | Fixed value |
| $\alpha$ | 0.24 | unitless | Fixed value |
| $\Theta$ | 0.85 | unitless | Fixed value |
| $R_{d,25}$ | | μmol m<sup>-2</sup> s<sup>-1</sup> | Gas exchange measurement |
| $\Delta H_a (R_d)$ | | J mol<sup>-1</sup> | Xue *et al.* (2017) |
| $\Delta H_a (K_o)$ | 23720 | J mol<sup>-1</sup> | |
| $c (K_o)$ | 14.68 | mmol mol<sup>-1</sup> | |
| $\Delta H_a (K_c)$ | 80990 | J mol<sup>-1</sup> | |
| $c (K_c)$ | 38.28 | μmol mol<sup>-1</sup> | Bernacchi *et al.* (2002) |
| $\Delta H_a (\Gamma^*)$ | 24460 | J mol<sup>-1</sup> | |
| $c (\Gamma^*)$ | 13.49 | μmol mol<sup>-1</sup> | |

Under well-watered conditions, the correlation between photosynthetic parameters and temperature can be expressed by the peak Arrhenius function (Dreyer, 2001; Medlyn *et al*., 2002; Xue *et al*., 2017; Kumarathunge *et al*., 2019; Knauer *et al*., 2019) as follows:

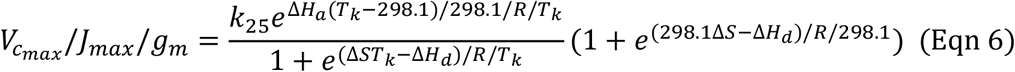

where *k*_25_ is the photosynthetic parameter value at 25C (*V*_cmax_,25, *J*_max,25_, or *g*_m,25_); Δ*H*_a_ is the activation energy; Δ*H*_d_ is the deactivation energy; Δ*S* is the entropy term which characterizes the changes in reaction rate caused by substrate concentration (Table 1); *T*_k_ is leaf temperature in Kelvin unit; and *R* is the gas constant (8.314 Pa m^3^ K^-1^ mol^-1^).

*V*_cmax_, *J*_max_, and *g*_m_ can be estimated from field measurements, whereas *C* remains unknown in the leaf photosynthesis model. The leaf photosynthesis model is therefore required to be coupled with the stomatal conductance sub-model for predicting the behavior of stomatal conductance by depending on environmental drivers and *A*n. The stomatal conductance sub-model can be expressed as follows:

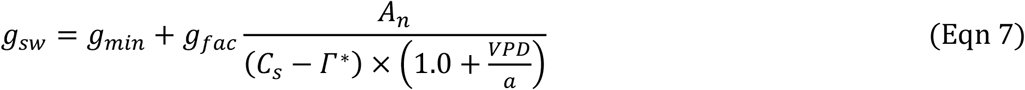

where *C*_s_ is leaf surface CO_2_ concentration; *a* is a constant (default as 35kPa); *g*_min_ is the value of *g*_sw_ when *A*_n_ is zero; *g*_fac_ is the stomatal sensitivity to the assimilation rate; and *VPD* is the vapor pressure deficit between the leaf surface and atmosphere; *g*_sw_ is the stomatal conductance to water vapor; and *g*_fac_ and *g*_min_ are the slope and intercept of the linear relationship between *g*_sw_ and *A*_n_, respectively, which were extracted from the diurnal gas exchange data.

To predict the response of *A*_n_ to leaf temperature at ambient and relatively high CO_2_ concentrations and light saturation levels (i.e. *A*_max_-*T*_leaf_ curve) for the species derived from literature, we needed to determine the *g*_fac_ and *g*_min_ values, without the diurnal gas exchange information. According to our field measurements, *g*_fac_ was set to 3.0 for the woody species and 4.0 for the herbaceous species. The *g*_min_ was set to 10.0 mmol m^-2^ s^-1^ for both C_3_ woody and herbaceous species, as commonly adopted by most TBMs (Sellers *et al*., 1996). An initial value was set for *C*, and then, *A*_n_ was determined using Eqn. 1. The known *A*_n_ was substituted into Eqn. 7 for *g*_sw_ estimation, which in turn was substituted into Eqn. 8 or 9 to generate a new *C* value. The new *C* value was then compared with the previous value. This new *C* value was adjusted and then considered as the second initial value for Eqn. 1 until the difference between the generated *C* and the previous one became less than 0.05 ppm. The iteration procedure used here (i.e. the Newton-Raphson iteration method) was consistent with that used by most TBMs (Dai *et al*., 2003; Sellers *et al*., 1996).

#### (1) The *g*_m_ infinite model and parameterization

In the *g*_m_ infinite model, the CO_2_ diffusion conductance from intercellular airspace to the chloroplast stroma was assumed to be infinitely large, a common practice in line with almost all TBMs. Photosynthesis is considered to be limited either by stomatal aperture/closure or by CO_2_ fixation, which depends on the functioning of leaf photochemistry and/or photosynthetic enzymes. *C* is the intercellular CO_2_ concentration. According to the Fick’s first law, *C* can be expressed as:

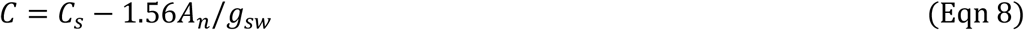

*V*_a_ was estimated using the linear phase of the *A*_n_/*C*_i_ curve (*C*_i_ from 50 to 200 ppm) and *J*_a_ was estimated using the saturated phase (*C*_i_ > 400 ppm) by referring to the studies by Xu and Badocchi (2003) and Kumarathunge *et al.* (2019). Other input parameters for the *g*_m_ infinite model were clarified in Methods S1.

#### (2) The *g*_m_ finite model and parameterization

In the *g*_m_ finite model, the total conductance of CO_2_ diffusion from the leaf surface to the chloroplast stroma consists of *g*_sw_ and *g*_m_. Photosynthesis is limited by three major factors: stomatal conductance, mesophyll conductance, and biochemical/photochemical limitations. *C* is the CO_2_ concentration inside the chloroplasts that depends on *A*_n_, *g*_sw_, *g*_m_, and *C*_s_. It can be expressed according to the Fick’s first law as follows:

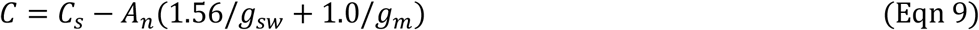

*V*_cmax_ and *J*_max_ are the maximum carboxylation rate and the maximum electron transport rate based on the CO_2_ concentration in the chloroplasts, respectively. In this study, *V*_cmax_, *J*_max_, and *g*_m_ values were estimated using four different parameter estimation methods. The Bayesian retrieval algorithm (Zhu *et al*., 2011; Han *et al*., 2020) and the Sharkey online calculator (Sharkey *et al*., 2007) were used to estimate *V*_cmax_, *J*_max_, and *g*_m_ values by using the *A*_n_/*C*_i_ curve only (abbreviated as *V*_cmax_B_, *J*_max_B_, *g*_m_B_ and *V*_cmax_S_, *J*_max_S_, and *g*_m_S_, respectively). The chlorophyll fluorescence-gas exchange method was used to estimates *g*_m_ by using the variable *J* method (abbreviated as *g*_m_F_), according to methods used by previous studies (Harley *et al*., 1992; Niinemets *et al*., 2009; Xue *et al*., 2016; 2017; Carriquí *et al*., 2020; 2021). The anatomy method for *g*_m_ estimation is constrained to a narrow range of leaf temperature around 25°C (*g*_m_A,25_) (Tosens *et al*., 2012; Tomas *et al*., 2013). In this study, the prior range of parameters for the Bayesian retrieval algorithm adopted the range recommended by Zhu *et al*. (2011) and Han *et al*. (2020). The prior ranges of *R*_d_ for woody plants and herb plants were 0.01-2.0 μmol m^-2^ s^-1^ and 0.01-5.0 μmol m^-2^ s^-1^, respectively. Notably, the unit of *g*_m_ estimated using the Sharkey online calculator and the Bayesian retrieval algorithm was μmol m^-2^ s^-1^ Pa^-1^; therefore, it was required to be converted into mol m^-2^ s^-1^ by using the formula: [(g_m_(mol m^-2^ s^-1^) = *g*_m_(μmol m^-2^ s^-1^ Pa^-1^) × P/10], where *P* is the actual atmospheric pressure (Pa). The explicit clarity on the parameter values assumed for each parameter estimation method was referred to the Methods S1.

*V*_cmax_, *J*_max_, and *g*_m_ estimated using the Sharkey online calculator, Bayesian retrieval algorithm, chlorophyll fluorescence-gas exchange method, and anatomy method were grouped to develop eight parameterization solutions to drive the *g*_m_ finite and infinite models (Table 2). For the plant species without *g*_m_A,25_ data, model parameterization solutions adopted *g*_m_B_, *g*_m_S_, and *g*_m_F_. *g*_m_FA_ at a leaf temperature of 25°C was the mean of *g*_m_F,25_ and *g*_m_A,25_. *g*_m_FA_ values at other temperatures were approximated by *g*_m_F_ only.

**Table 2.**
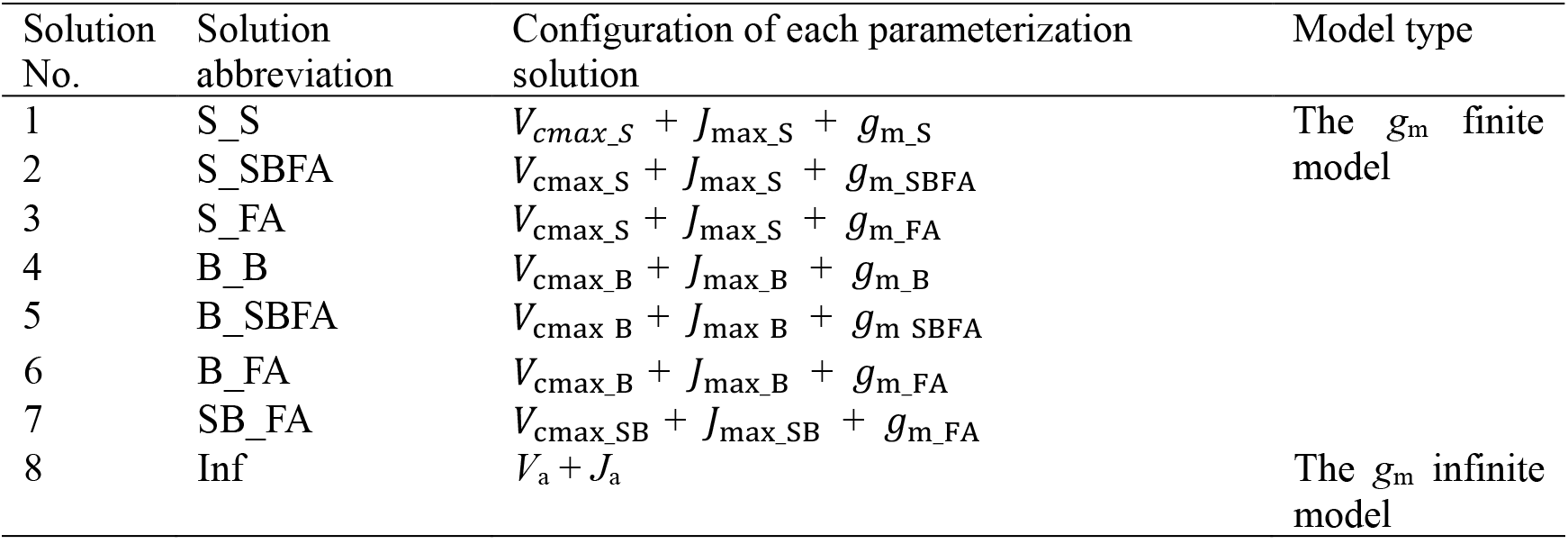
Eight parameterization solutions considered to drive the *g*_m_ infinite and finite photosynthesis-transpiration coupled models. *V*_a_: the apparent maximum carboxylation rate; *J*_a_: the apparent maximum electron transport rate; *V*_cmax_B_: *V*_cmax_ estimated using the Bayesian retrieval algorithm; *V*_cmax_S_: *V*_cmax_ estimated using the Sharkey online calculator; *V*_cmax_SB_: the mean of *V*_cmax_B_ and *V*_cmax_S_; *J*_max_B_: *J*_max_ estimated using the Bayesian retrieval algorithm; *J*_max_S_: *J*_max_ estimated using the Sharkey online calculator; *J*_max_SB_: the mean of *J*_max_B_ and *J*_max_S_; *g*_m_B_: *g*_m_ estimated using the Bayesian retrieval algorithm; *g*_m_S_: *g*_m_ estimated using the Sharkey online calculator; *g*_m_F_: *g*_m_ estimated using the chlorophyll fluorescence-gas exchange method; *g*_m_A,25_: *g*_m_ estimated using the anatomy method; *g*_m_SBFA_: the mean of *g*_m_S_, *g*_m_B_, *g*_m_F_ and *g*_m_A,25_; and *g*_m_FA_: the mean of *g*_m_F_ and *g*_m_A,25_.

Variation in the Rubisco kinetic parameters measured *in vitro* was less than 10% amongst C_3_ plant species (von Caemmerer, 2020); thus these parameters may be assumed to be identical for all vegetation types. In this study, the use of the Rubisco kinetic parameters (*K*_c_, *K*_o_, and *Γ*\*) for the *g*_m_ finite model was consistent with Bernacchi *et al*. (2002) (Table 1). However, Knauer *et al*. (2019; 2020) used two types of Rubisco kinetic parameters for model comparisons, of which one type was adopted from the study by Bernacchi *et al*. (2002) for the *g*_m_ finite model and the other type was adopted from a study by Bernacchi *et al*. (2001) for the *g*_m_ infinite model. Similarly, the Rubisco kinetic parameters in study of Bernacchi *et al*. (2001) were used for parameterization of the *g*_m_ infinite model in 141 C_3_ species by Kumarathunge *et al*. (2019). The hypotheses proposed by Knauer *et al*. (2019; 2020) and Kumarathunge *et al*. (2019) were evaluated in this study. Hence, it is necessary here to parameterize the *g*_m_ infinite model by using the 2001 version in association with the Rubisco kinetic parameters.

### Data collection for model validation

*V*_cmax_, *J*_max_, and *g*_m_ estimations by the four parameter estimation methods were performed using field measurements of the *A*_n_/*C*_i_ curve plus chlorophyll fluorescence at leaf temperatures ranging from 10-15°C to 40°C in 19 species under 31 experimental treatments (four tropical deciduous tree species, four deciduous broadleaf tree species, seven evergreen broadleaf tree species, three C_3_ crops species, and one C_3_ herb and grass species) (Methods S2 and S3). Diurnal gas exchange rates were measured in 15 species under 25 treatments (Methods S2 and S3). Leaves were sampled *in-situ* immediately after the gas exchange measurement to determine the leaf anatomical structure (Methods S4). For parameter correlation analysis, the *A*_n_/*C*_i_ curve and chlorophyll fluorescence data at 25°C in seven gymnosperms specie, five ferns species and four herbs species were collected from literature (Carriquí *et al*., 2020; Nadal *et al*., 2018) (Methods S5). For the *T*_optA_-*T*_opt_gm_ correlation analysis, data on *T*_optA_ and optimum temperature of *g*_m_ (*T*_opt_gm_) estimated using the carbon isotope discrimination method (*g*_m__^13^c) and the chlorophyll fluorescence-gas exchange method in five C_3_ crops species, two C_3_ herbs and grasses species, and one deciduous broadleaf tree species were collected from literature (Evans and von Caemmerer, 2013; Li *et al*., 2020; Scafaro *et al*., 2011; von Caemmerer and Evans, 2015; Warren and Dreyer, 2006; Xue *et al*., 2016) (Methods S6). Abbreviations of sampled species under different experimental treatments were referred to Methods S2-S6.

### Evaluations of the *g*_m_ infinite and finite models

Independent field data, including the *A*_max_-*T*_leaf_ curve, *T*_optA_, and diurnal gas exchange, were used to validate the *g*_m_ infinite and finite models. We fitted the *A*_max_-*T*_leaf_ curve in Eqn. 10 to obtain *T*_optA_ (Sall and Pettersson, 1994; Battaglia *et al*., 1996; Gunderson *et al*., 2009; Kumarathunge *et al*., 2019),

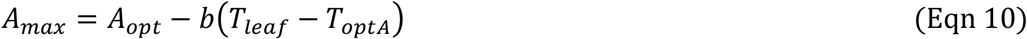

where *A*_opt_ is the *A*_n_ at *T*_optA_, and parameter *b* (unitless) describes the curvature of *A*_max_ and *T*_leaf_.

Root mean square error (RMSE) and Nash-Sutcliffe efficiency (NSE) coefficients were used to quantify the performance of the model (Methods S7).

## Results

### Temperature responses of *V*_cmax_, *J*_max_, and *g*_m_

The temperature curve-fitting lines for *V*_cmax_B_, *V*_cmax_S_, *V*_cmax_SB_, and *V*_a_ increased rapidly with increase in *T*_leaf_ from 10C to 35C, and began to decrease after reaching a peak (about 300 μmol m^-2^ s^-1^ for *V*_cmax_B_, *V*_cmax_S_, and *V*_cmax_SB_; 180 μmol m^-2^ s^-1^ for *V*_a_) at a leaf temperature around 35C. *V*_cmax_B_, *V*_cmax_S_, and *V*_cmax_SB_ presented a similar behavior *(p* > 0.1, Fig. 1a and Figs. S1a-ae), except at high leaf temperatures ≥ 35°C. Despite *V*_cmax_SB_ was significantly higher by 43.9% in average than *V*_a_ at 25°C, considerable scatter in the difference for the two variables was observed (Table S1). There were similar values at each leaf temperature among temperature response curves of *J*_max_B_, *J*_max_S_, *J*_max_SB_, and *J*_a_ (Fig. 1b), with the difference among them being less than 6% at 25°C and the optimal temperatures changing around 32-34°C (Figs. S2a-ae and Table S2). The *g*_m_B_ temperature curve-fitting line was overlapped with that of *g*_m_S_ (Fig. 1c), whereas both had significantly higher values by 69% and 58% than that of *g*_m_FA_ at 25°C, respectively (Table S3). However, the *g*_m_S_ temperature curve-fitting line did not well overlap with the 95% confidence interval because of irregular observations in *g*_m_S_ temperature response among sampled species/treatments (Figs. S3a-ae). The temperature curve-fitting lines for *g*_m_B_, *g*_m_S_, and *g*_m_FA_ exhibited larger variations in *T*_opt_gm_, ranging from 27 to 32°C, which were lower by 15.7% and 10.6% in average as compared to optimal temperatures for *V*_cmax_ (i.e. *V*_cmax_B_, *V*_cmax_S_, and *V*_cmax_SB_) and *J*_max_ (i.e. *J*_max_B_, *J*_max_S_, *J*_max_SB_), respectively.

**Fig. 1.**
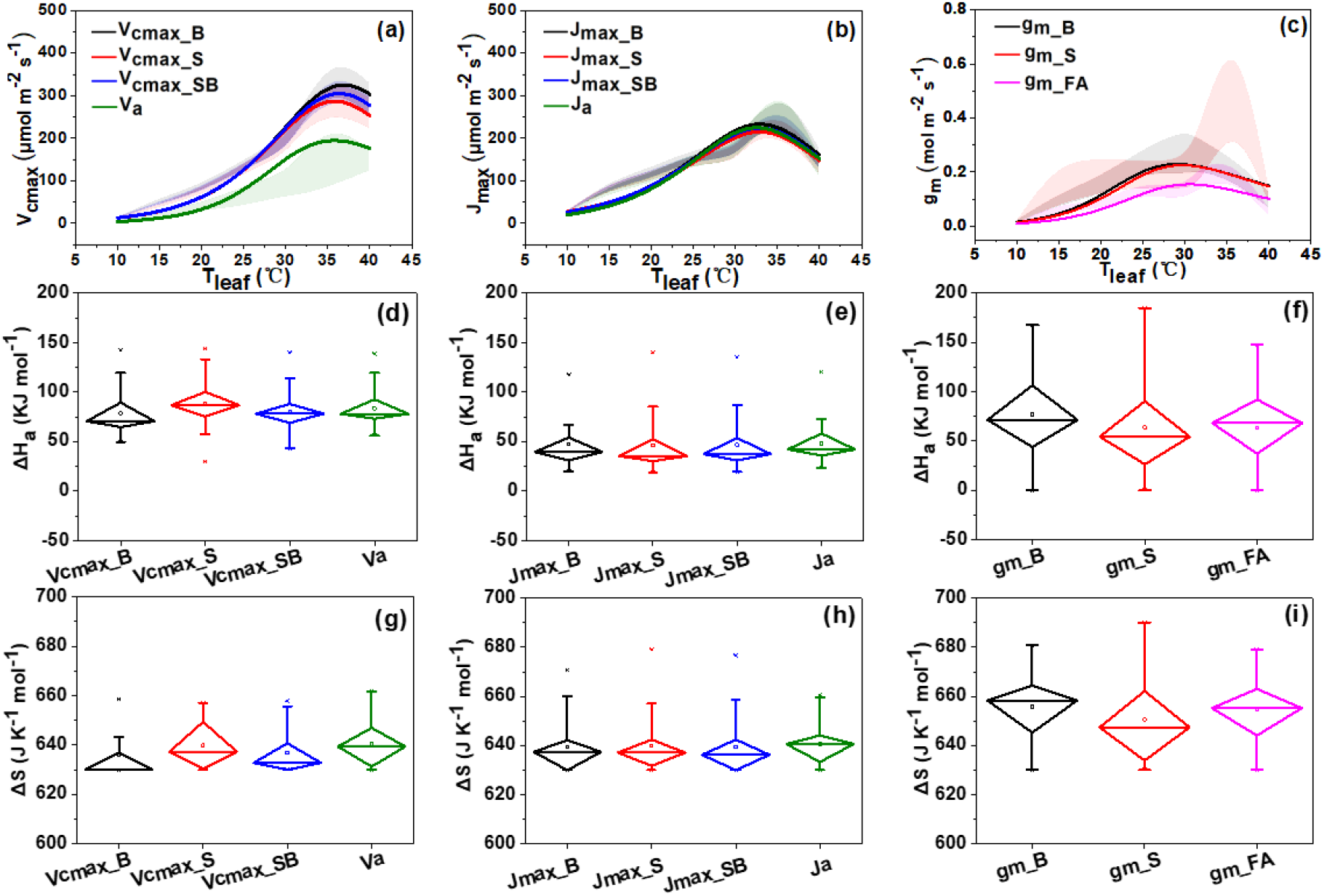
*V*_a_ and *V*_cmax_, *J*_a_ and *J*_max_, and *g*_m_ temperature response curves performed by fitting the mean of parameter estimations of 19 species at 31 treatments using the Eqn. 6 and were labeled by shadow zones (Figs. 1a-c). The shadow zone for each curve-fitting line was the 95% confidence interval of parameter estimations of all sampled species/treatments. Distribution features of Δ*H*_a_ and *ΔS* values that determine *V*_a_ and *V*_cmax_, *J*_a_ and *J*_max_, and *g*_m_ temperature response curves in all sampled species/treatments were displayed using the box-plot (Figs. 1d-i). The curve-fitting lines of *V*_a_ and *V*_cmax_, *J*_a_ and *J*_max_, and *g*_m_ temperature response for individual species/treatment were shown in Supporting Information Figs. S1-S3. *V*_cmax_, *J*_max_, and *g*_m_ values were estimated using different parameter estimation methods, namely the Bayesian retrieval algorithm (*V*_cmax_B_, *J*_max_B_, and *g*_m_B_, black solid line), the Sharkey online electronic calculator (*V*_cmax_S_, *J*_max_S_, and *g*_m_S_, red solid line), the mean of *V*_cmax_B_ and *V*_cmax_S_ and the mean of *J*_max_B_, and *J*_max_S_ (*V*_cmax_SB_ and *J*_max_SB_, blue solid line), the mean of *g*_m_ estimated using the chlorophyll fluorescence-gas exchange and anatomy methods (*g*_m_FA_, pink solid line), and the apparent maximum carboxylation rate and the apparent maximum electron transport rate (*V*_a_ and *J*_a_, green solid line).

The boxplot minimum value of Δ*H*_a_ was 49.2 KJ mol^-1^ for *V*_cmax_B_, 52.3 KJ mol^-1^ for *V*_cmax_S_, 37.6 KJ mol^-1^ for *V*_cmax_SB_, and 65.6 KJ mol^-1^ for *V*_a_ (Fig. 1d). For *J*_max_B_, *J*_max_S_, *J*_max_SB_, and *J*_a_, their minimum values were 22.2, 20.8, 22.6, and 23.6 KJ mol^1^, respectively (Fig. 1e). Similarities in the minimum value that was close to zero were observed for *g*_m_B_, *g*_m_s_, and *g*_m_FA_ (Fig. 1f). Zero value in Δ*H*_a_ was found in Pop_WC for *g*_m_B_, in Pop_CW for *g*_m_s_, and in E_sal and E_mdel for *g*_m_FA_ (Table S3 and Figs. S3u, t, q, and r). There were similarities in the interquartile range (IQR) of Δ*H*a among *V*_cmax_B_, *V*_cmax_S_, *V*_cmax_SB_, and *V*_a_, so did the Δ*H*_a_ of the four *J*_max_ types and the Δ*H*a of the three *g*_m_ types. However, larger ranges in the IQR of Δ*H*_a_ for *g*_m_ (i.e. *g*_m_B_, *g*_m_S_, and *g*_m_FA_) than those of *J*_max_ and *V*_cmax_ were clearly observed and coefficient of variation *(CV)* in Δ*H*a of *g*_m_ (i.e. *g*_m_B_, *g*_m_S_, and *g*_m_FA_) was amplified in average by 135.2% (Table S3). The IQR of Δ*S* for *V*_cmax_B_, *V*_cmax_S_, *V*_cmax_SB_, and *V*_a_ was 5.1, 20.2, 8.3, and 15.6 J K^-1^ mol^-1^, respectively (Fig. 1g). Similarities in the IQR of *ΔS* among *J*_max_B_, *J*_max_S_, *J*_max_SB_, and *J*_a_ temperature responses were evident (Fig. 1h and Table S2). Whereas, the first quartile (Q1) values of Δ*S* for *g*_m_ (i.e. *g*_m_B_, *g*_m_s_, and *g*_m_FA_) were similar or slightly higher than the third quartile (Q3) values of *V*_cmax_ (i.e. *V*_cmax_B_, *V*_cmax_S_, and *V*_cmax_SB_) and *J*_max_ (i.e. *J*_max_B_, *J*_max_S_, and *J*_max_SB_) (i.e. mean of the Q1 value for *g*_m_ was 644.5 J K^-1^ mol^-1^; mean of the Q3 value for *V*_cmax_ was 644.1 J K^-1^ mol^-1^, and mean of the Q3 value for *J*_max_ was 647.5 J K^-1^ mol^-1^). The mean value of Δ*S* across *g*_m_B_, *g*_m_S_, and *g*_m_FA_ temperature curves was 654 J K^-1^ mol^-1^, whereas it was 639 J K^-1^ mol^-1^ for *J*_max_ (i.e. *J*_max_B_, *J*_max_S_, and *J*_max_SB_) and 637 J K^-1^ mol^-1^ for *V*_cmax_ (i.e. *V*_cmax_B_, *V*_cmax_S_, and *V*_cmax_SB_) (Table S1-S3). The *CV*s of Δ*S* for *g*_m_B_, *g*_m_S_, and *g*_m_FA_ temperature curves were 1.74%, 2.88%, and 2.2%, respectively, while the corresponding *CV*s of Δ*H*_a_ were 58.45%, 78.98%, and 53.26%, respectively (Table S3). These results suggested: greater variations in Δ*H*_a_ and Δ*S* of *g*_m_ than those of *V*_cmax_ and *J*_max_; large variations in Δ*H*_a_ and relatively small variations in Δ*S* for *g*_m_; and greater values in Δ*S* for *g*_m_ than those for *V*_cmax_ and *J*_max_.

### Comparisons between gas exchange observations and simulations by the *g*_m_ finite and infinite models

For the *A*_max_-*T*_leaf_ curves measured at ambient *C*_a_ = 400-450 ppm, the slopes of the linear regression between observations and simulations by the S_S, S_SBFA, and S_FA parameterization schemes were 0.87, 0.91, and 1.00, respectively (Figs. 2a-c). The adjusted R^2^ (adj.R^2^) values for the three schemes were 0.81, 0.86, and 0.90, respectively, and the corresponding NSE values were 0.79, 0.84, and 0.88. The RMSE values of the three parameterization schemes accounted for 24%, 22%, and 19% of the mean *A*_max_ (*A*_max,mean_ = 12.47 μmol m^-2^ s^-1^). The predicted *A*max values from the S_S and S_SBFA schemes were 11% and 7% higher than the *A*_max,mean_, respectively (Figs. S4a-ae). These results suggested that *g*_m_ estimation using the chlorophyll fluorescence-gas exchange method and anatomy method was more reasonable. In line with the results obtained from S_S, S_SBFA, and S_FA schemes, the B_FA scheme fitted *A*_max_ at each temperature better than the B_B and B_SBFA schemes (Figs. 2d-f and Figs. S4a-ae), which further suggested that the configuration of *g*_m_FA_ for *g*_m_ estimation was more reasonable. As shown in Figs. 1a and b, the values of *V*_cmax_B_ and *V*_cmax_S_ and those of *J*_max_B_ and *J*_max_S_ at each temperature were similar to each other. Therefore, we considered the possible effects of the combination of *V*_cmax_ or *J*_max_ estimations using the Bayesian retrieval algorithm and Sharkey online calculator on *A*_max_ simulations (the SB_FA scheme in Figs. S4a-ae). The NSE value for the SB_FA scheme (0.92) was 5% and 1% higher, whereas the ratio of RMSE to *A*_max,mean_ was 21% and 12% lower than those for the S_FA and B_FA schemes suggesting better *A*_max_ predictions using the parameter configuration scheme by considering *V*_cmax_SB_, *J*_max_SB_, and *g*_m_FA_. The adj.R^2^, NSE, and the ratio of RMSE to *A*_max,mean_ were 0.85, 0.73, and 28%, respectively (Fig. 2h), indicating that although the *g*_m_ infinite model is generally credible in estimating the *A*_max_-*T*_leaf_ curve, the prediction errors in this model are relatively larger. We found that *A*_max_ predictions using the *g*_m_ infinite model were significantly higher (16%) than the observations (partially for 30-40°C, Figs. S4a-ae). Additionally, numerical simulation indicated that the *g*_m_ finite model driven by the *g*_m_ temperature function without considering the deactivation stage significantly overestimated *A*_max_ at higher temperatures 30-35-40°C by 10.99%, 32.51%, 64.06%, respectively. The results obtained from the eight parameterization schemes at *C*_a_ = 600-630 ppm were similar to those at *C*_a_ = 400-450 ppm (Figs. 2a-h and Figs. S5a-ae). These results indicated that the *g*_m_ finite model driven by the SB_FA scheme could predict *A*max more accurately than the *g*_m_ infinite model.

**Fig. 2.**
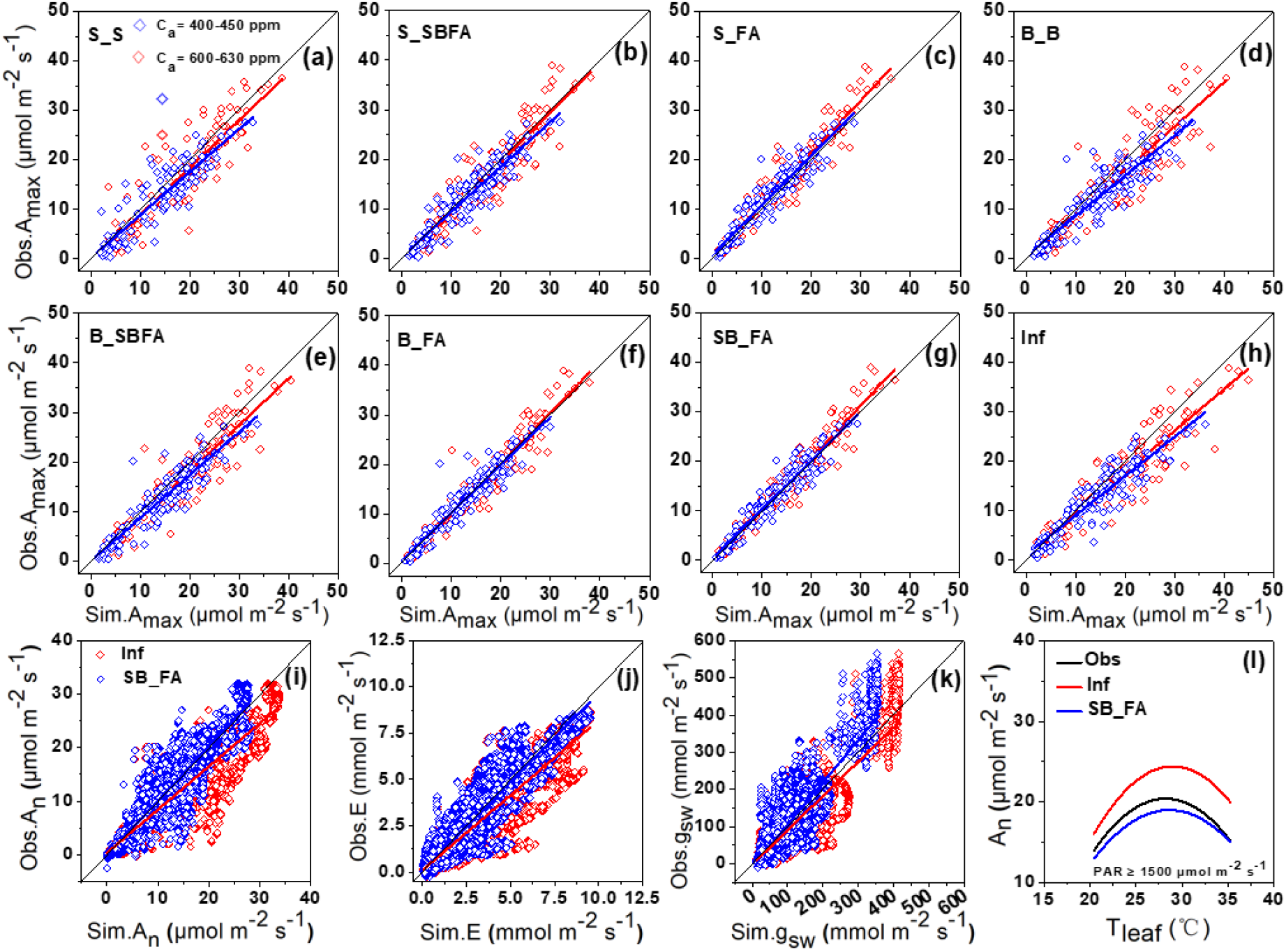
Scatter plots showing comparisons in *A*_max_ observations and predictions between the *g*_m_ infinite and finite models that are forced by eight parameterization schemes (Figs. 2a-h, S_S, S_SBFA, S_FA, B_B, B_SBFA, B_FA, SB_FA, and Inf). The blue solid line signifies the linear regression fit between observations at *C*_a_ = 400-450 ppm and predictions, and the red solid line signifies the linear regression fit between observations at *C*_a_ = 600-630 ppm and predictions. Scatter diagrams in Figs. 2i-k showing comparisons in the diurnal assimilation rate (*A*_n_), transpiration rate (*E*), and stomatal conductance (*g*_sw_) between observations and predictions using the *g*_m_ infinite model (red diamonds) and the *g*_m_ finite model that is forced by the parameterization scheme SB_FA (blue diamonds). The black solid line represents the 1:1 line. Curve-fitting lines in Fig. 2l show comparisons in diurnal *A*_n_ at *PAR* greater than 1500 μmol m^-2^ s^-1^ and across a wide range of leaf temperatures between observations (black line) and predictions using the *g*_m_ infinite model (red line) and the *g*_m_ finite model that is forced by the parameterization scheme SB_FA (blue line). *C*_a_: CO_2_ concentration at leaf surface; *PAR:* photosynthetically active radiation.

Figs. 2i-l shows the comparisons in diurnal *A*_n_, *E,* and *g*_sw_ between observations and predictions using the *g*_m_ infinite model and the *g*_m_ finite model that is forced by the parameterization scheme SB_FA (detailed comparisons in each species/treatment shown in Figs. S6-S8). The adj.R^2^ values for *A*_n_, *E*, and *g*_sw_ predicted using the *g*_m_ finite model were 0.88, 0.85, and 0.78, respectively, whereas those predicted using the *g*_m_ infinite model were 0.87, 0.83, and 0.76, respectively. The NSE values for the three variables for the *g*_m_ finite model were 0.88, 0.85, and 0.76, respectively, and those for the *g*_m_ infinite model were 0.76, 0.75, and 0.74, respectively. The ratio of RMSE to the mean of *A*_n_, *E*, and *g*_sw_ for the *g*_m_ finite model was 27%, 34%, and 40%, respectively, whereas that for the *g*_m_ infinite model was 38%, 45%, and 42%, respectively. Meanwhile, significant overestimations in simulated *A*_n_ under heat shocking conditions (i.e. *T*_leaf_ from > 30°C) by 25%-40% by using the *g*_m_ infinite model were observed (Fig. 2l and Fig. S9). These results suggested that the *g*_m_ finite model is superior to the *g*_m_ infinite model in predicting diurnal gas exchange under a wide range of growth conditions.

For the diurnal changes in *A*_n_, the adj.R^2^ and NSE for the *g*_m_ finite model were improved by 1% and 12% compared with those for the *g*_m_ infinite model. For the diurnal changes in *E*, the adj.R^2^ and NSE for the *g*_m_ finite model improved by 2% and 10% compared with those for the *g*_m_ infinite model. These results suggested that the effects of *g*_m_ finite expression on photosynthesis and transpiration estimations were almost equally stronger.

### Statistical correlations between *T*_optA_ and photosynthetic parameters

The linear regression slope and adj.R^2^ values between *T*_optA_ predictions using the *g*_m_ finite model and observations were 0.67 and 0.85, respectively, whereas those obtained using the *g*_m_ infinite model were 0.88 and 0.18, respectively (Fig. 3a). *T*_optA_ predicted using the *g*_m_ finite model displayed a better correlation with the observations. We observed that *T*_optA_ observations had a significant positive correlation with the *T*_opt_gm_ derived from *g*_m_FA_ and *g*_m__^13^C temperature responses, and the adj.R^2^ value reached 0.58 (Fig. 3b). No significant correlations were found among *T*_optA_-Δ*H*_a_ of *g*_m_FA_, *T*_optA_-*g*_m_FA,25_, and *T*_optA_-*J*_max_SB,25_/*V*_cmax_SB,25_ (Fig. S10).

**Fig. 3.**
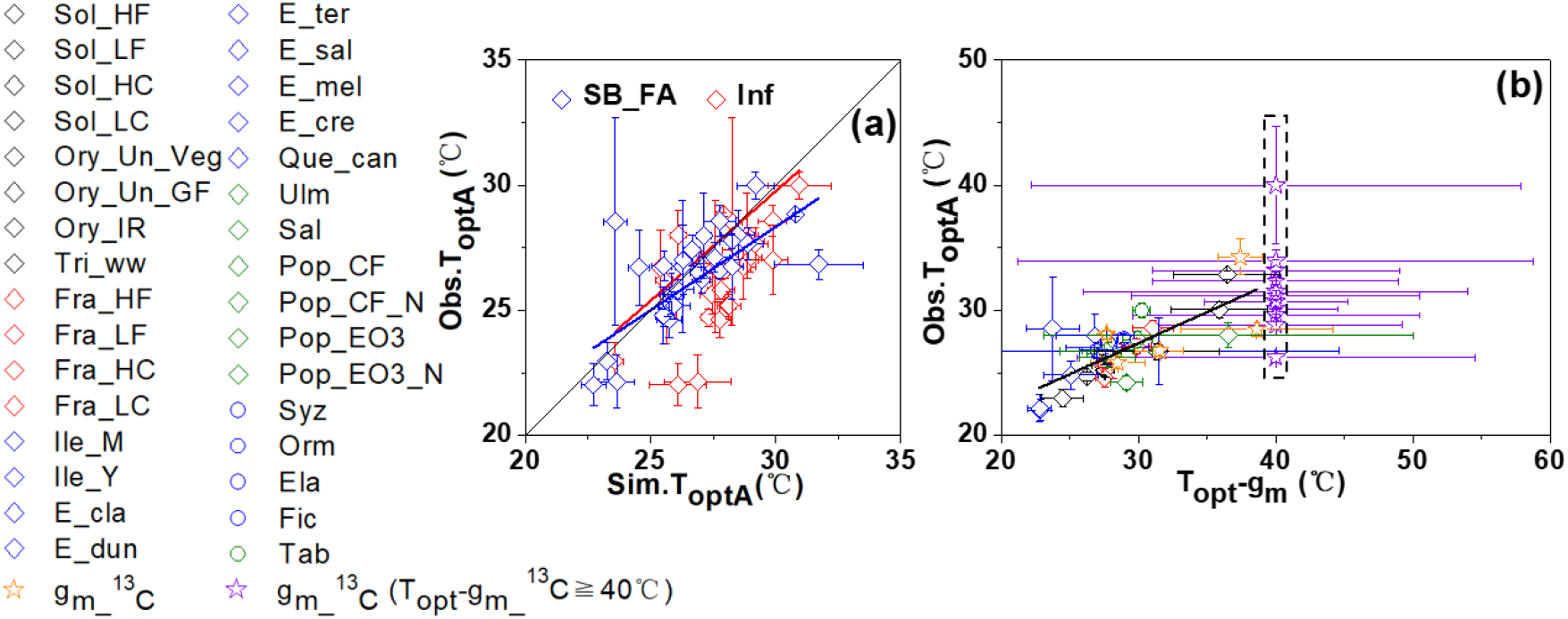
Scatter diagrams showing comparisons in photosynthesis optimal temperature (*T*_optA_) between observations (Obs.*T*_optA_) and predictions using the *g*_m_ finite model forced by the parameterization solution SB_FA (Sim. *T*_optA_, blue circles) and by the *g*_m_ infinite model (red circles) (a) and the statistical correlation for Obs. *T*_optA_ and *g*_m_ optimal temperature (*T*opt-gm) across different plant functional types (b). The black solid line in Fig. 3a represents the 1:1 line. The red and blue lines in Fig. 3a and the black line in Fig. 3b are the linear regression fit. *T*_opt-gm_ in *g*_m_ temperature response estimated using the ^13^C discrimination method greater than and equal to 40°C (*T*_opt-gm__^13^C values ≥ 40°C were unified to be 40°C indicated by purple stars inside the dashed square) was not included for statistical correlation analysis in Fig. 3b. The sampled species/treatments are classified into six plant functional types (PFTs): black diamonds for C_3_ crops (C3C), blue diamonds for evergreen broadleaf trees (EBF), green diamonds for deciduous broadleaf trees (DBF), red diamonds for C_3_ herbs and grasses (C3G), blue circles for tropical evergreen broadleaf trees (TRF), and green circles for tropical deciduous broadleaf trees (TDF).

### Statistical correlations among photosynthetic parameters estimated by four parameter estimation methods

*g*_m_F,25_ was positively correlated with *g*_m_A,25_, whereas the adj.R^2^ was 0.22, and the linear regression slope was 0.57, which is much lower than 1.0 (Fig. 4a), primarily because of large discrepancies between the two variables in rice plants. The two variables strongly correlated in other sampled species/treatments with the adj.R^2^ of 0.76 and the slope of 0.80. A significant exponential correlation was found between *g*_m_FA,25_ and *g*_m_SB,25_ (adj.R^2^ = 0.67, *p* < 0.01, Fig. 4b), so did the correlation between *g*_m_F,40_ and *g*_m_SB,40_ (<u>gray circles in Fig. 4b</u>), with a close agreement found at low *g*_m_ < 0.15 mol m^-2^ s^-1^. A negative correlation for *g*_m_FA,25_ and the difference between *V*_cmax_SB,25_ and *V*_a,25_ was found (adj.R^2^ = 0.47,*p* < 0.05, Fig. 4c). *J*_max_SB,25_ was closely related to *V*_cmax_SB,25_ (adj.R^2^ = 0.83,*p* < 0.01, Fig. 4d).

**Fig. 4.**
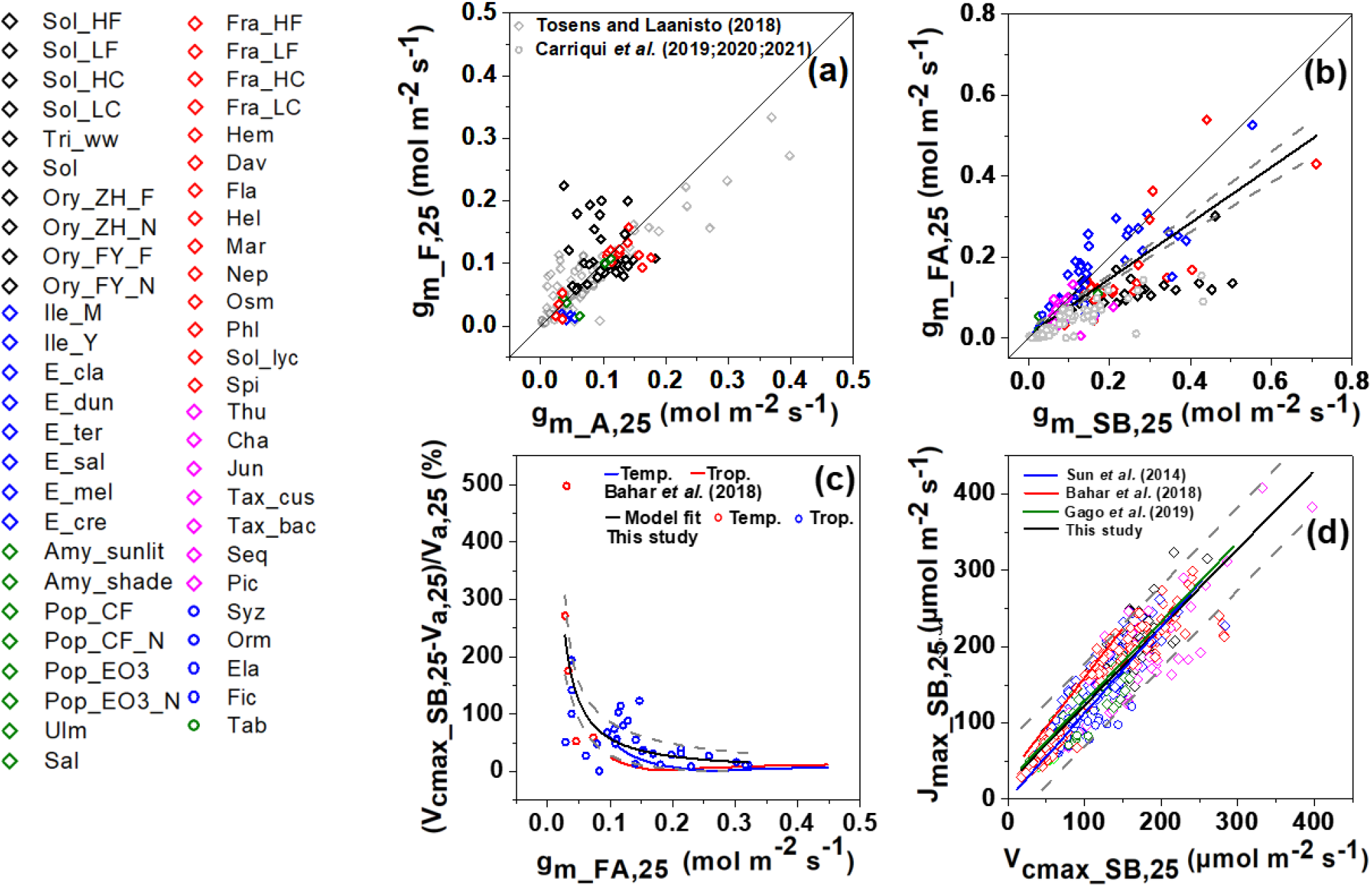
Statistical correlations for *g*_m_F,25_-*g*_m_A,25_ (a), for *g*_m_FA,25_-*g*_m_sB,25_ (b), for *g*_m_FA,25_-the difference between *V*_cmax_SB,25_ and *V*_a,25_ (c), and for *J*_max_SB,25_ *V*_cmax_SB,25_ (d) in sampled species/treatments. The sampled species/treatments in Figs. 4a, b and d are classified into six plant functional types (PFTs): black diamonds for C_3_ crops (C3C), blue diamonds for evergreen broadleaf trees (EBF), green diamonds for deciduous broadleaf trees (DBF), red diamonds for C_3_ herbs and grasses (C3G), blue circles for tropical evergreen broadleaf trees (TRF), and green circles for tropical deciduous broadleaf trees (TDF). The paired datasets for *g*_m_F,25_ and *g*_m_A,25_ in studies of Tosens and Laanisto (2018) and Carriquí *et al.* (2019; 2020; 2021) were added in Fig. 4a (grey diamonds and circles). *g*_m_SB,25_ greater than 1.5 mol m^-2^ s^-1^ are regarded as outliers and are not shown. Gly_amb, Pop_WC, and Pop_CW are not shown as they do not have the observation data at a leaf temperature of 25°C. *g*_m_F,40_ and *g*_m_SB,40_ datasets were added (gray circles) in Fig. 4b for comparison with the *g*_m_FA,25_-*g*_m_SB,25_ correlation. *V*_cmax_Cc_ and *V*_cmax_Ci_ values in temperate species (<u>Temp</u>.: *Phyllocladus aspleniifolius)* and tropical species (<u>Trop</u>.: *Litsea leefeana)* in the study of Bahar *et al.* (2018) were used to calculate the difference of the two variables as shown in Fig. 4c (blue and red lines, respectively). *V*_cmax_Cc_ in their study (i.e. *V*_cmax,25_) was derived from the *A*_n_/*C*_c_ curve that is converted from the *A*_n_/*C*_i_ curve by assuming a constant *g*_m_ at the entire *C*_i_ range. *V*_cmax_Ci_ is the apparent *V*_cmax_ (i.e. *V*_a,25_) derived from the *A*_n_/*C*_i_ curve in their study, which is equivalent to *V*_a_ in our study. The *g*_m,25_ in their study was estimated using the ^13^C isotope discrimination method. The dashed lines in grey color represent the 95% prediction intervals of the nonlinear regression. <u>Temp.: temperate species; Trop.: tropical species</u>.

## Discussion

### Superiority of the *g*_m_ finite model and an optimized parameterization solution that excludes the equifinality phenomenon

The *A*_max_-*T*_leaf_ curves were accurately predicted using the *g*_m_ infinite model only in 15 species/treatments (Figs. S4d, j-s, y, aa, ad, and ae), whereas *A*_max_ values at higher temperatures (30-40°C) were largely overestimated in other sampled species/treatments. Poor modeling accuracies were also evident in the simulation of diurnal gas exchange rates (Fig. 2l and Figs. S6c, e, h, k, l, n, p, and q). It aligned with the previous reports in rice and winter wheat by Xue *et al*. (2016) and in *Quercus ilex* by Niinemets *et al*. (2009) who found that the photosynthesis model that does not explicitly consider *g*_m_ cannot always predict leaf gas exchange accurately. The *g*_m_ finite model considers the dependent effects of chloroplast CO_2_ concentration on the assimilation rate, stomatal and mesophyll resistance, and leaf surface CO_2_ concentration (von Caemmerer, 2013; 2020). The relationships among CO_2_ diffusion flux, CO_2_ concentration gradient between the leaf surface and chloroplasts, and gas diffusion resistance may be approximated using the Fick’s first law (Harley *et al*., 1992; Xue *et al*., 2017), as shown in Eqn. 9. Simulation results in gas exchange in 19 species under a wide range of growth conditions proved that the *g*_m_ finite model has good transferability and high prediction capacity, due to which the model can be applied to as many C_3_ species as possible. The *g*_m_ finite model developed in this study can be conveniently substituted into most TBMs by simply replacing (*C*_i_ = *C*_s_ – 1.56*A*_n_/*g*_sw_) as given in Eqn. 9.

*V*_cmax_, *J*_max_, and *g*_m_ temperature responses are key photosynthetic parameters of the *g*_m_ finite model. Despite a “one-point” method has been proposed to quantify *V*_cmax_, there is no consensus that this method can be widely used (Burnett *et al*. 2019). One way to determine *V*_cmax_ is from the *A*_n_/*C*_c_ curve that is converted from the *A*_n_/*C*_i_ curve using a value for *g*_m_ determined at ambient CO_2_ and assuming a constant *g*_m_ for the entire range of *C*_i_ (Manter and Kerrigan, 2004; Bahar *et al*., 2018). Whereas, the conversion of *A*_n_/*C*_i_ to *A*_n_/*C*_c_ may not be true at low *g*_m_ values under certain conditions (Flexas *et al*., 2007). The key photosynthetic parameters can also be quantified using the curve-fitting methods applied to the *A*_n_/*C*_i_ curve; for example, the Bayesian retrieval algorithm and the Sharkey online calculator (Sharkey *et al*., 2007; von Caemmerer*et al*., 2009; Gu *et al*., 2010; Zhu *et al*., 2011; von Caemmerer, 2013). The curve-fitting methods that simultaneously quantify the three key photosynthetic parameters may have the equifinality phenomenon (i.e., similar simulation results can be achieved with different initial conditions), because of a curvilinear negative correlation between *V*_cmax_ estimation and *g*_m_ (von Caemmerer, 2013) that changes steeply at the lower range of *g*_m_ (generally < 0.1 mol m^-2^ s^-1^) (Bahar *et al*., 2018). For the parameterization of the *g*_m_ finite model, *V*_cmax_, *J*_max_, and *g*_m_ values were simultaneously estimated by the Bayesian retrieval algorithm, so did the Sharkey online calculator. Importantly, *g*_m_ was also determined using the chlorophyll fluorescence-gas exchange method and the anatomy method. Our results indicated that the *g*_m_B_ temperature responses were significantly different from the *g*_m_S_ temperature responses in terms of absolute values at 25°C and Δ*H*_a_ in most sampled species/treatments (Figs. S3a-b, d, f-i, l, o-t, y, aa-ac), whereas similarities in the temperature response for either *V*_cmax_ or *J*_max_ were found between the two curve-fitting methods. It implied that the differences in *V*_cmax_ observed at high temperatures between the two methods (Fig. 1a) were likely not related to *g*_m_ estimation. A close agreement between the *g*_m_F,25_ and *g*_m_A,25_ (Fig. 4a) and between *g*_m_FA,25_/40 and *g*_m_SB,25_/40, especially at the lower range < 0.15 mol m^-2^ s^-1^ (Fig. 4b), was evident, respectively. Furthermore, there is a close agreement in *T*_opt_gm_ between the chlorophyll fluorescence-gas exchange method and the Bayesian retrieval method (adj.R^2^ = 0.58 and the regression slope = 0.81, Fig. S3a-ae). Results suggested that the four parameter estimation methods give similar *g*_m_ estimations, especially at the lower range. The consequence of variations in *g*_m_ on estimation of *V*_cmax_ on *C*c and *C*_i_ bases in this study was consistent with the reports by Bahar *et al.* (2018) (Fig. 4c). Hence, the estimations in *V*_cmax_ by fusing the Bayesian retrieval algorithm and the Sharkey online calculator are reasonable. Furthermore, we found a significantly linear correlation between *V*_cmax_SB,25_ and *J*_max_SB,25_ (Fig. 4d), which falls between the reports by Bahar *et al.* (2018) and Sun *et al.* (2014) and overlaps with the reports by Gago *et al*. (2019). These results suggested that the four parameter estimation methods are independent in the measurements taken for parameter estimation and optimization algorithms, namely the optimized parameterization scheme (i.e. the SB_FA scheme, discussed below) is physiologically meaningful, which excludes the equifinality phenomenon.

In the present study, a close relationship between *g*_m_F,25_ and *g*_m_A,25_ was observed in most sampled species/treatments (Fig. 4a). The close correspondence between the two variables has been also reported in many other C_3_ species (Tosens and Laanisto, 2018; Carriquí *et al*., 2019; 2020; 2021) (grey symbols shown in Fig. 4a). Mesophyll conductance is a complex three-dimensional trait that is probably determined by both biochemical and anatomical features. The chlorophyll fluorescence method compares gas exchange signal with the optical signal which may vary with the depth through the mesophyll tissue (Evans, 2021). These variations may explain minor discrepancies between *g*_m_A,25_ and *g*_m_F,25_ in most sampled species/treatments (Fig. 4a). Carriquí *et al*. (2020) reported that the cell wall composition is a key factor in the *g*_m_ setting in sclerophyll species. The constant values in model parameters of the anatomy *g*_m_ estimation method for vascular plants such as the ratio of cell wall porosity to tortuosity (*P*_cw_) may also contribute to the discrepancies. In this study, *g*_m_F,25_ was found to be significantly higher than *g*_m_A,25_ in rice. An extremely dense distribution of mesophyll cells in rice was observed (ultrastructure images not shown). The total length of the chloroplast facing the intercellular space (*l*_c_) in association with *g*_m_A,25_ by using the ultrathin sections was probably underestimated in rice because of the exclusion of tightly adjacent parts of two mesophyll cells in the sampling fields of view.

As shown in Figs. 2a and d, *g*_m_ estimation using the *A*_n_/*C*_i_ curve only through the Bayesian retrieval algorithm or the Sharkey online calculator can accurately predict leaf gas exchange rates in some species/treatments, whereas cannot in others. A better modeling performance was obtained by the S_FA scheme (i.e. *V*_cmax_S_+*J*_max_S_+*g*_m_FA_) than the S_S scheme (i.e. *V*_cmax_S_+*J*_max_S_+*g*_m_s_). Our results are similar to reports by Sharkey *et al*. (2007): if chlorophyll fluorescence data are available, it could be possible to estimate *g*_m_ from those data to ameliorate reliability of modeling performance. Results suggested the SB_FA solution (i.e. *V*_cmax_SB_+*J*_max_SB_+*g*_m_FA_) that can predict the photosynthesis and transpiration better than other parameterization solutions as the optimized parameterization solution for the *g*_m_ finite model.

### Temperature responses characteristics of *g*_m_

A *g*_m_ response to temperature has been reported in some species, but not in others (Scafaro *et al*., 2011; von Caemmerer and Evans, 2015; Shrestha *et al*., 2019; Evans, 2021; Li *et al*., 2020), partially due to differences in *g*_m_ estimation methods used by them, as seen in great variations in *g*_m_ temperature response obtained by different estimation methods (Fig. 1c and Figs. S3a-ae). We argued that the *g*_m_ estimated using the parameter estimation method that can accurately fit the *A*_max_-*T*_leaf_ curve and diurnal gas exchange rates has higher credibility than those that cannot predict accurately. Results of our study suggested that 90% of the sampled species under well-watered conditions exhibit a significant response of *g*_m_ to temperature. Inter-species variations in both Δ*H*a and Δ*S* for *g*_m_ were significantly greater than those of *V*_cmax_ and *J*_max_. Large variations in Δ*H*a signify significant changes in *g*_m_ temperature response across species, which is in agreement with the results of the studies by Shrestha *et al.* (2019), von Caemmerer and Evans (2015), and Evans (2021). The two-components modeling method developed by von Caemmerer and Evans (2015) produced Δ*H*_a_ for CO_2_ permeability through the membranes, ranging between 36 and 76 kJ mol^-1^. Using the isolated pea leaf plasma membranes, Zhao *et al.* (2017) reported Δ*H*_a_ for CO_2_ permeability of 30.2 and 52.4 kJ mol^-1^ at high and low internal carbonic anhydrase (CA) concentrations. Results of our study reported the lower and upper limits of the 95% confidence interval of Δ*H*_a_ for *g*_m_FA_ were 50.14 and 76.12 kJ mol^-1^, respectively. The determined Δ*H*_a_ values of our study are similar to the ranges reported by von Caemmerer and Evans (2015) and Zhao *et al.* (2017). The rates of CO_2_ diffusion in the membranes and during the liquid phase reflect the amount of CO_2_-permeable and transport enzyme proteins, their thermal stabilities, and the structural components of cell wall. <u>A common or distinct set of a</u>quaporins in the membranes and <u>the associating proteins such as</u> CA inside the vesicles that affect Δ*H*_a_ for CO_2_ permeability/diffusion (Zhao *et al*., 2017) likely vary greatly in expression levels and the associating heterotetramers among C_3_ species (Otto *et al*., 2010; Momayyezi *et al*., 2020). Hence, the determined large variations in temperature response attributes of *g*_m_ are reasonable.

The temperatures at which *V*_cmax_ and *J*_max_ deactivate were usually higher than 35°C and even 40°C in most sampled species. They are similar to the findings in tobacco (Bernacchi*et al*., 2001), rice (Xue *et al*., 2016), and poplar (Silim *et al*., 2010; Xu *et al*., 2020). We found that *g*_m_ deactivation temperatures were lower than those for *V*_cmax_ and *J*_max_ (Figs. 1a-c), which are similar to reports by Xu *et al*. (2020) and Warren and Dreyer (2006). Meanwhile, better accuracy in leaf gas exchange predictions was obtained using the temperature peak function for *g*_m_ than the monotone increasing function. Results highlighted importance of incorporating the deactivation stage of *g*_m_ into leaf photosynthesis model for better modeling accuracy. A rapid change in fluidity of plasma membrane was observed within 1-3 min after heat shock of 37°C in *A. thaliana* and wheat (Zheng *et al*., 2012; Abdelrahman *et al*., 2020). <u>Elevated temperature increases membrane permeability. During a 40°C, 5-15 min heat stress Triandafillou *et al.* (2020) found that *Saccharomyces cerevisiae* cells rapidly acidify from a pH of 7.5 to a range of slightly acidic pH values around 6.8 due to proton influx, which could constrain the CA stability by 20-30% (Wang *et al*., 2016). The transient intracellular acidification induced by heat stress is broadly conserved in eukaryotes</u>. Changes in fluidity and permeability of plasma membrane at elevated temperatures (Niu and Xiang, 2018) that cause proton influx and impact the CA stability may cause the decline of *g*_m_ by 10-20% at elevated temperature from 30°C to 40°C (Fig. 1c).

### Effects of the *g*_m_ finite expression on photosynthesis and transpiration

Kumarathunge *et al*. (2019) proposed that the variation in *T*_optA_ among species can be explained by biochemical restrictions (*J*_a,25_/*V*_a,25_). *V*_a_ and *J*_a_ are referred to as the apparent maximum carboxylation rate and electron transport rate, respectively, and both these parameters, particularly *V*_a_, are numerically and physiologically different from *V*_cmax_ and *J*_max_. A similar result of an asymmetric effect of *g*_m_ estimation on *V*_cmax_ and *J*_max_ estimations was reported by Manter and Kerrigan (2004) and Sun *et al*. (2014). Hence, we may argue that *V*_a_ not only represents the Rubisco carboxylation rate but also embeds the *g*_m_ effects. Stripping off the *g*_m_ effects from *V*_a,25_ would cause changes in *J*_a,25_/*V*_a,25_ and then cause significant changes in the *T*_optA_-*J*_a,25_/*V*_a,25_ correlation. Our results suggested that *T*_optA_ has a significant linear correlation with *T*_opt_gm_ but not *JV*r, Δ*H*_a_ of *g*_m_FA_, and *g*_m_FA,25_, implying that *T*_optA_ is probably related to *g*_m_ temperature response characteristics. Whereas, the plasticity in *T*_optA_ was not fully related to *g*_m_ because only 58% of the variations in *T*_optA_ were explained by *g*_m_. Changes in the Rubisco kinetic properties, such as thermal stability of Rubisco activase (Salvucci and Crafts-Brandner, 2004), could be an important mechanism.

Knauer *et al.* (2020) argued that the *g*_m_ finite expression has significant effects on photosynthesis estimation and that the effects of *g*_m_ on transpiration are marginal. Conversely, we found that the effects of the *g*_m_ finite expression on photosynthesis and transpiration simulations are equally stronger. Significant effects on transpiration estimation were probably achieved through better predictions of *g*_sw_ because transpiration is a product of *VPD* and *g*_sw_. Across diverse species, *g*_m_ is strongly linked with *g*_sw_ and leaf hydraulic conductance through the *g*_m_ linkage to extra-xylem components (Flexas *et al*., 2013) such as the plasma membrane intrinsic proteins (PIPs) subfamily of aquaporins (Groszmann *et al*., 2017). Physiological mechanisms underlying the integrated hydraulic-photosynthetic system explained the observed effects of the *g*_m_ finite expression on photosynthesis and transpiration.

## Conclusions

In this study, we developed a *g*_m_ finite photosynthesis-transpiration coupled model that can be directly applicable for most TBMs and also proposed an optimized parameterization solution. The *g*_m_ finite model driven by the parameterization theme of *V*_cmax_SB_, *J*_max_SB_ and *g*_m_FA_ could well predict *A*_max_-*T*_leaf_ curves and diurnal gas exchange rates in all sampled species under various experimental treatments. However, the *g*_m_ infinite model cannot always accurately track variations in photosynthesis and transpiration. Results suggested large variations in Δ*H*_a_ and Δ*S* for *g*_m_. *T*_optA_ was related to thermal attributes of *g*_m_ not *JV*r. Meanwhile, the explicit *g*_m_ expression had equally important effects on photosynthesis and transpiration estimations at plant species level. Results of our study proved that the *g*_m_ finite expression in most TBMs is important for better understanding effects of *g*_m_ on photosynthesis and transpiration under climate change.

## Acknowledgements

This research was supported by the Fundamental Research Funds for the Central Universities (lzujbky-2020-28) and the National Natural Science Foundation of China (32001129). Sincere thanks are dedicated to Dr. John R. Evans for his comments on the manuscript. We are also grateful to Dandan Liu and Xiao Song for their helps on field data collection. The authors declare that they have no conflicts of interest.

## Author contributions

WX: design of the research and funding, data analysis, collection and interpretation, and manuscript writing and revision. HL: data analysis, collection and interpretation, and manuscript writing. JE, MC, MN, TH, and CW: data collection and analysis, manuscript revision. J-F H, J-L Z, Z-G Y and X-W F: data collection and part work of data analysis.

## Data availability

The data that support the findings of this study are available from the corresponding author upon reasonable request. MATLAB script for Bayesian retrieval algorithm and the *g*_m_ finite photosynthesis-transpiration coupled model compiled by FORTRAN can be obtained through directly contacting the correspondence author.

## Supporting Information

Additional supporting information may be found in the online version of this article: **Fig. S1** The temperature response curves of *V*_cmax_.

**Fig. S2** The temperature response curves of *J*_max_.

**Fig. S3** The temperature response curves of *g*_m_.

**Fig. S4** Temperature response curves of the net assimilation rate under ambient CO_2_ concentration (400-450 ppm) and high radiation derived from field measurements and predictions.

**Fig. S5** Temperature response curves of the net assimilation rate under ambient CO_2_ concentration (600-630 ppm) and high radiation derived from field measurements and predictions.

**Fig. S6** Comparisons in diurnal photosynthesis rate between field observations and predictions.

**Fig. S7** Comparisons in diurnal transpiration rate between field observations and predictions.

**Fig. S8** Comparisons in diurnal stomatal conductance between field observations and predictions.

**Fig. S9** Comparisons in diurnal *An* at PAR greater than 1500 μmol m^-2^ s^-1^ and across a wide range of leaf temperatures between observations and predictions using the *gm* infinite model and the *g*_m_ finite model that is forced by the parameterization scheme SB_FA.

**Fig. S10** Statistical relationships for photosynthesis optimal temperature and the ratio of the maximum carboxylation rate to the maximum electron transport rate (*JV*r) (a), *T*_optA_-*g*_m_F,25_ (b), and *T*_optA_--activation term of *g*_m_F_ (Δ*H*_a_) (c).

**Table S1** Temperature response characteristic parameters of *V*_cmax_.

**Table S2** Temperature response characteristic parameters of *J*_max_.

**Table S3** Temperature response characteristic parameters of *g*_m_.

**Methods S1** The explicit clarity on the parameter values assumed for each parameter estimation method

**Methods S2**

**Methods S3**

**Methods S4**

**Methods S5**

**Methods S6**

**Methods S7**

