## Supplemental Files for "Explicit expression of mesophyll conductance in the traditional leaf photosynthesis–transpiration coupled model and its physiological significances"

New Phytologist

***New Phytologist* Supporting Information**

Article acceptance date: Click here to enter a date.

The following Supporting Information is available for this article:

**Fig. S1** Temperature response curves of the maximum carboxylation rate estimated using the Bayesian retrieval algorithm ($V_{cmax\_B}$, black solid line), the Sharkey online electronic calculator ($V_{cmax\_S}$, red solid line), the mean of $V_{cmax\_B}$ and $V_{cmax\_S}$ ($V_{cmax\_SB}$, blue solid line) and the apparent maximum carboxylation rate ($V_a$, black dashed line) in 19 species under 31 experimental conditions.

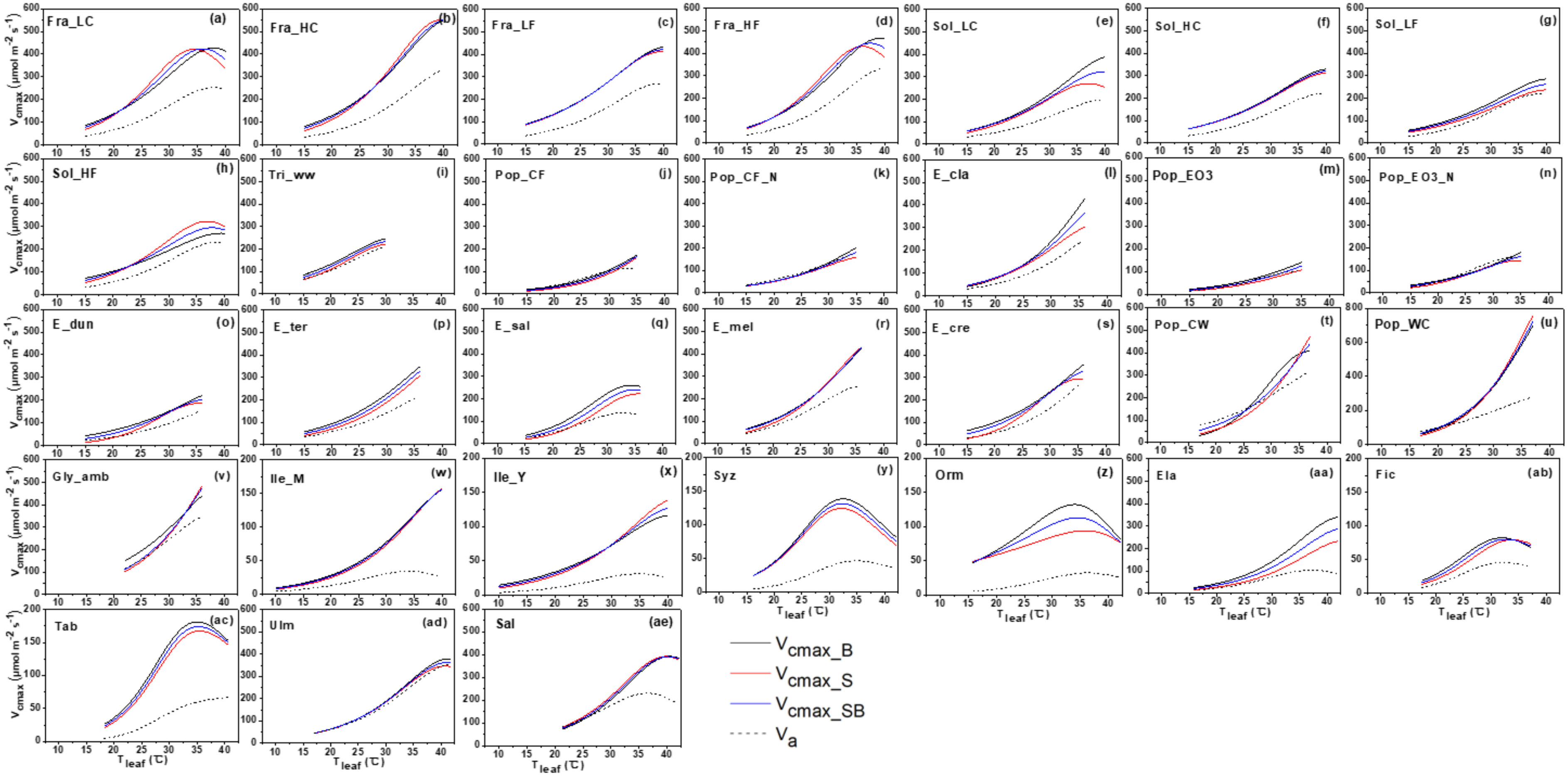

New Phytologist

**Fig. S2** Temperature response curves of the maximum electron transport rate estimated by Bayesian retrieval algorithm ($J_{max\_B}$, black solid line), the Sharkey online electronic calculator ($J_{max\_S}$, red solid line), the mean of $J_{max\_B}$ and $J_{max\_S}$ ($J_{max\_SB}$, blue solid line) and the apparent maximum electron transport rate ($J_a$, black dashed line) in 19 species under 31 experimental treatments.

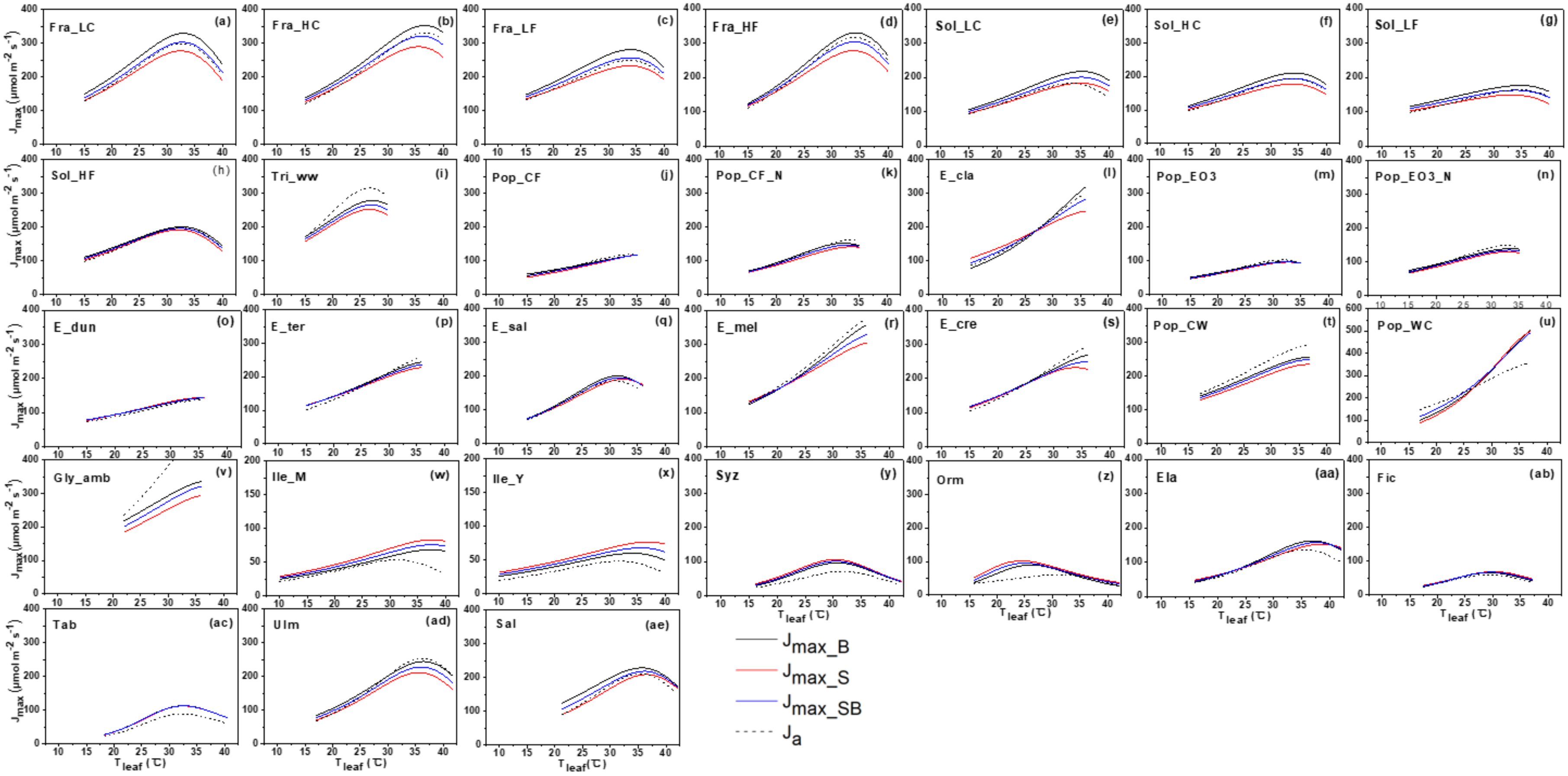

**Fig. S3** Temperature response curves of mesophyll conductance estimated by Bayesian retrieval algorithm ($g_{m\_B}$, black solid line), the Sharkey online electronic calculator ($g_{m\_S}$, red solid line) and the mean value of $g_m$ values estimated by gas exchange-chlorophyll fluorescence and anatomical methods ($g_{m\_FA}$, blue solid line) in 20 species under 31 experimental treatments. $g_{m\_B}$ and $g_{m\_S}$ use the left axis, and $g_{m\_FA}$ uses the right axis.

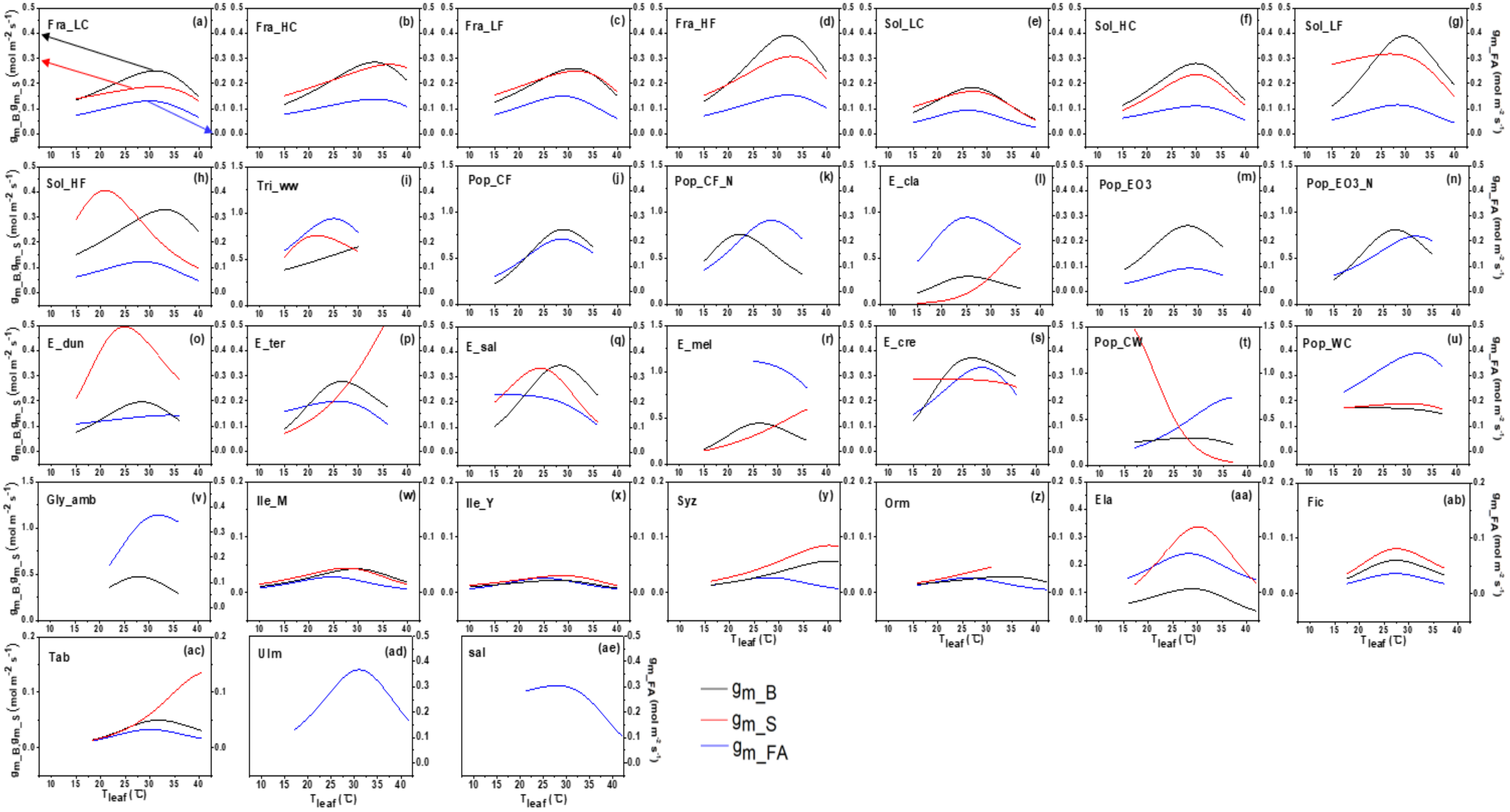

**Fig. S4** Temperature response curves of the net assimilation rate at saturating light conditions under an ambient $CO_2$ concentration (400-450 ppm) derived from field measurements and predicted by the $g_m$ infinite and finite models, which in turn are driven by eight parameterization schemes. Black diamond symbols indicate estimations from field data, the blue dotted line represents predictions by the parameterization scheme S_S (Table 2), the blue dashed line represents predictions by the parameterization scheme S_SBFA, the blue solid line for predictions by the parameterization scheme S_FA, the green dotted line for predictions by the parameterization scheme B_B, the green dashed line for predictions by the parameterization scheme B_SBFA, the green solid line for predictions by the parameterization scheme B_FA, the black solid line for predictions by the parameterization scheme SB_FA, the black dashed line for predictions by the parameterization scheme Inf.

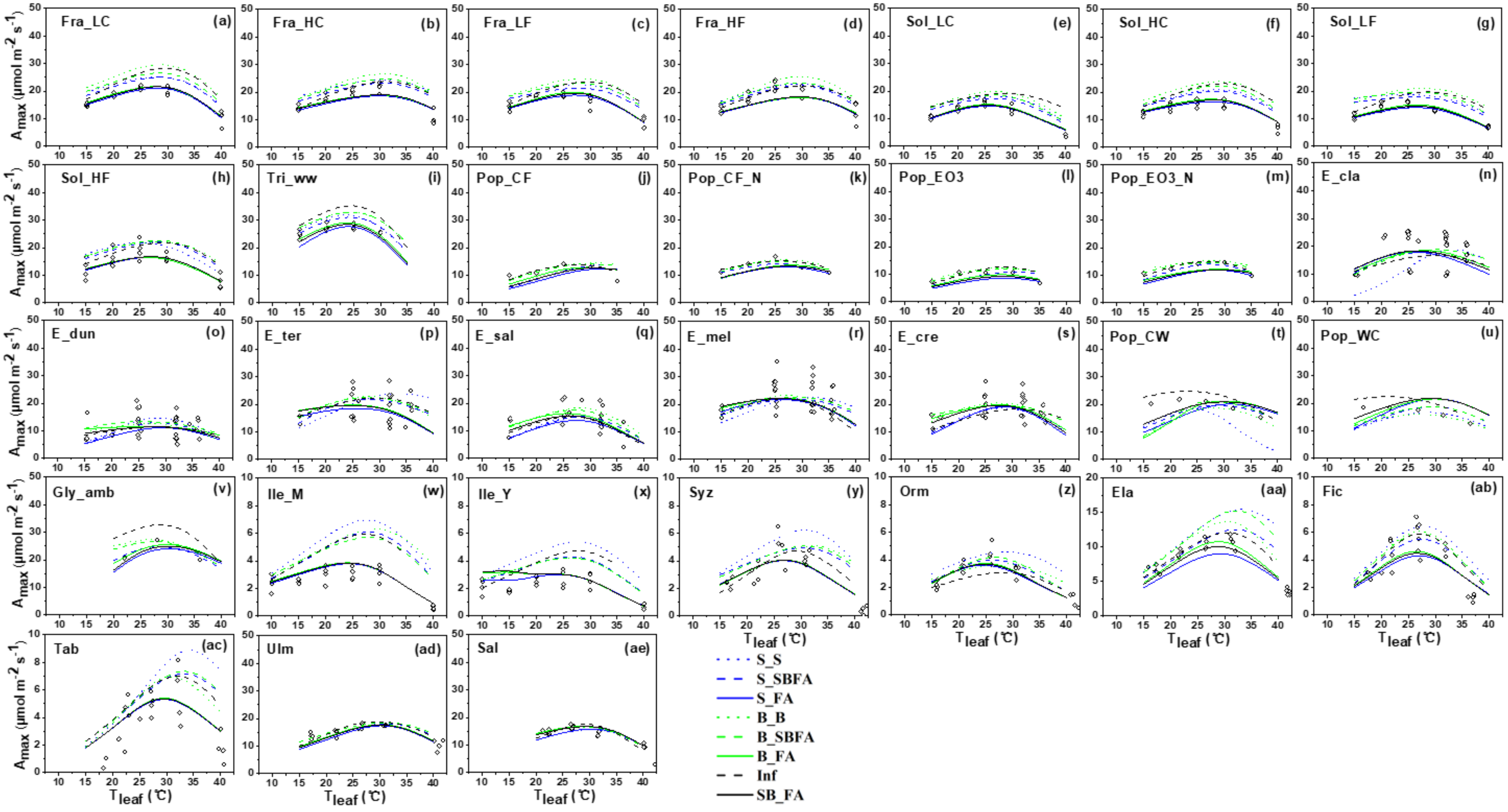

**Fig. S5** Temperature response curves of the net assimilation rate under ambient $CO_2$ concentration (600-650 ppm) and high radiation derived from field measurements and predictions by the $g_m$ infinite and finite models that are parameterized by eight parameterization schemes. The black diamond indicates field observations, the blue dotted line for predictions by the parameterization scheme S_S, the blue dashed line for predictions by the parameterization scheme S_SBFA, the blue solid line for predictions by the parameterization scheme S_FA, the green dotted line for predictions by the parameterization scheme B_B, the green dashed line for predictions by the parameterization scheme B_SBFA, the green solid line for predictions by the parameterization scheme B_FA, the black solid line for predictions by the parameterization scheme SB_FA, the black dashed line for predictions by the parameterization scheme Inf.

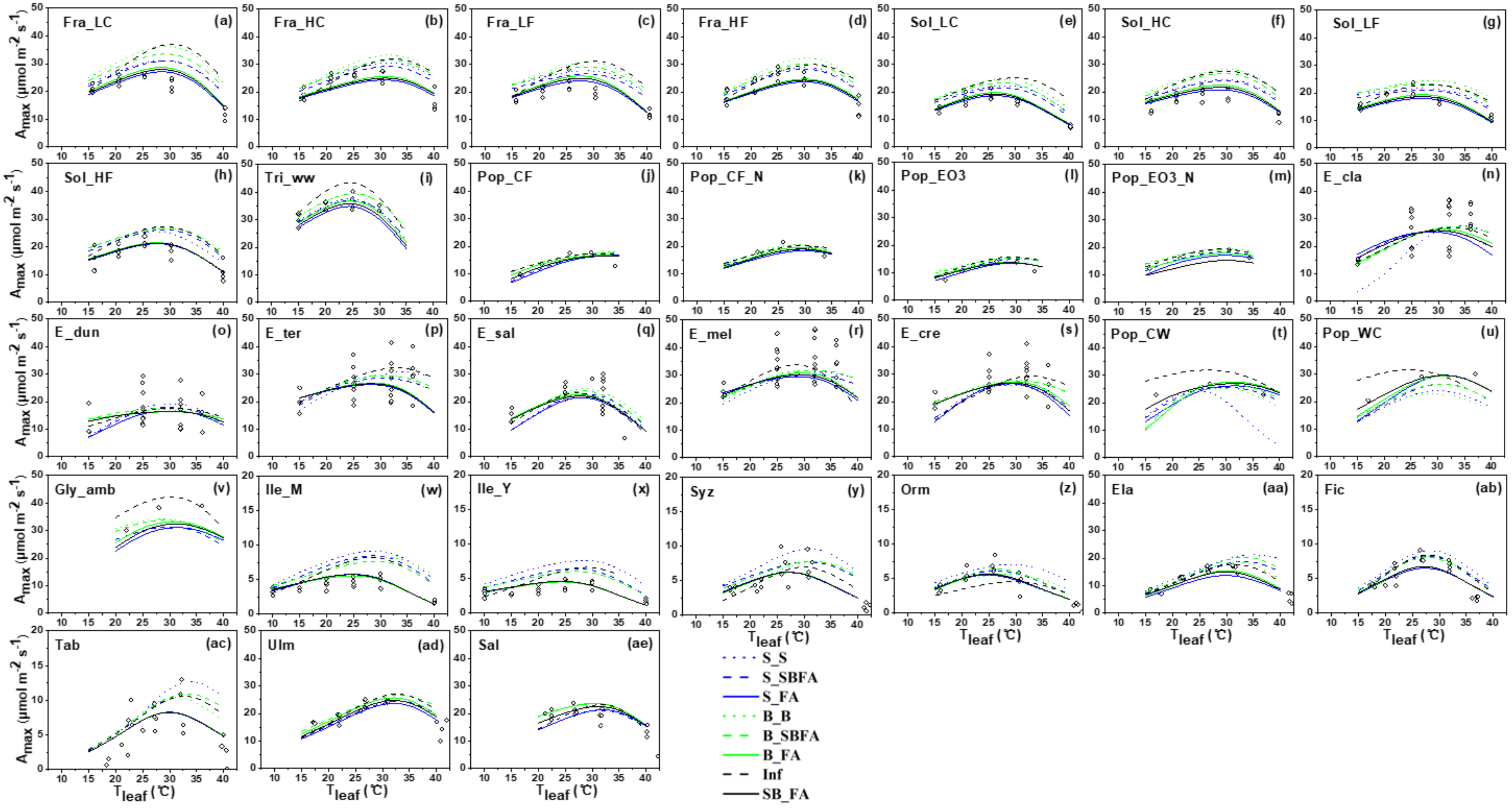

**Fig. S6** Comparisons in diurnal photosynthesis rate ($A_n$) between field observations (black diamonds) and predictions by the $g_m$ finite model driven by the parameterized scheme SB_SBFA (blue lines) and the $g_m$ infinite model driven by the parameterized scheme Inf (red lines)

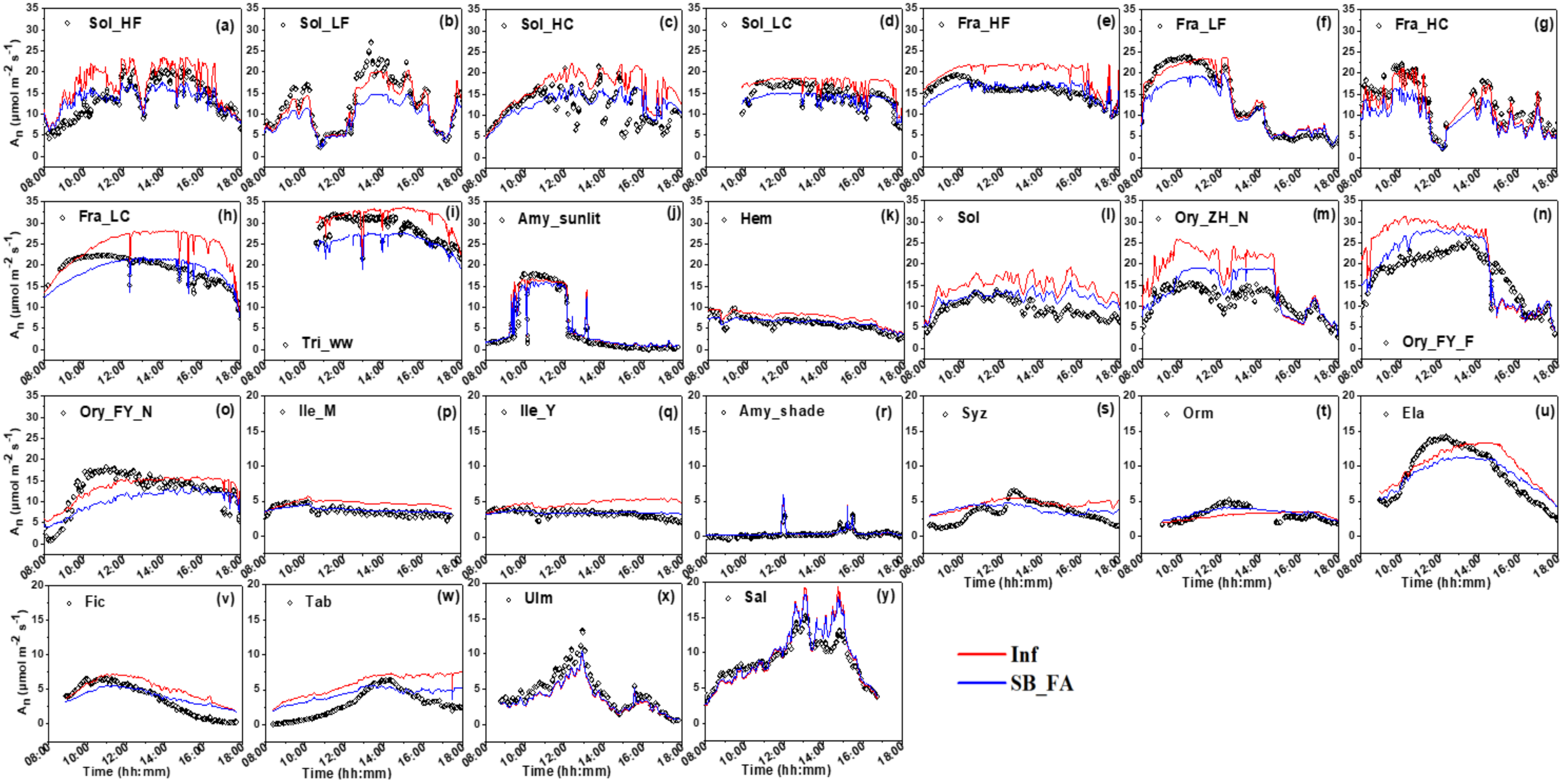

**Fig. S7** Comparisons in diurnal transpiration rate (*E*) between field observations (black diamonds) and predictions by the $g_m$ finite model driven by the parameterized scheme SB_SBFA (blue lines) and the $g_m$ infinite model driven by the parameterized scheme Inf (red lines)

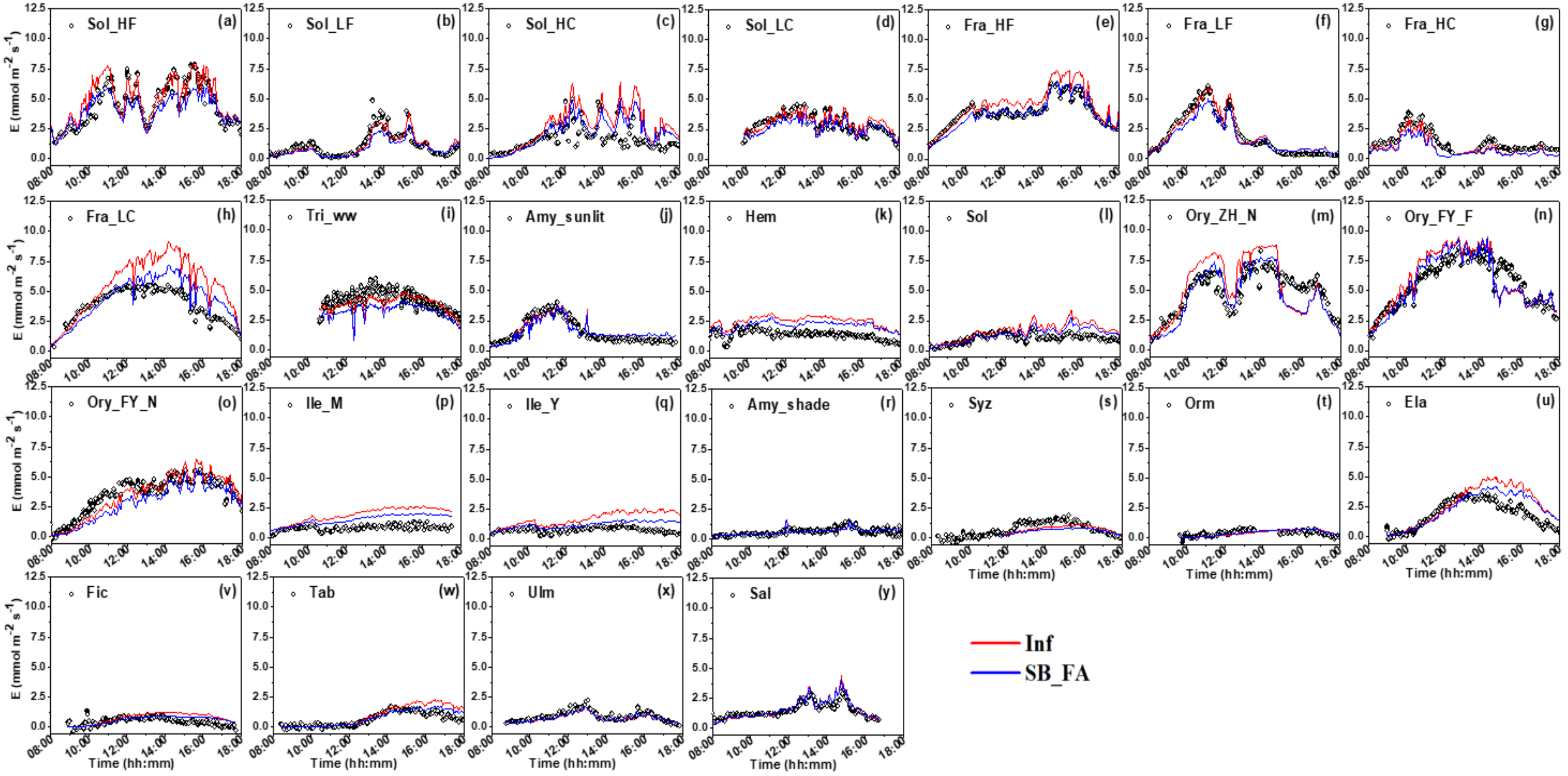

**Fig. S8** Comparisons in diurnal stomatal conductance ($g_{sw}$) between field observations (black diamonds) and predictions by the $g_m$ finite model driven by the parameterized scheme SB_SBFA (blue lines) and the $g_m$ infinite model driven by the parameterized scheme Inf (red lines).

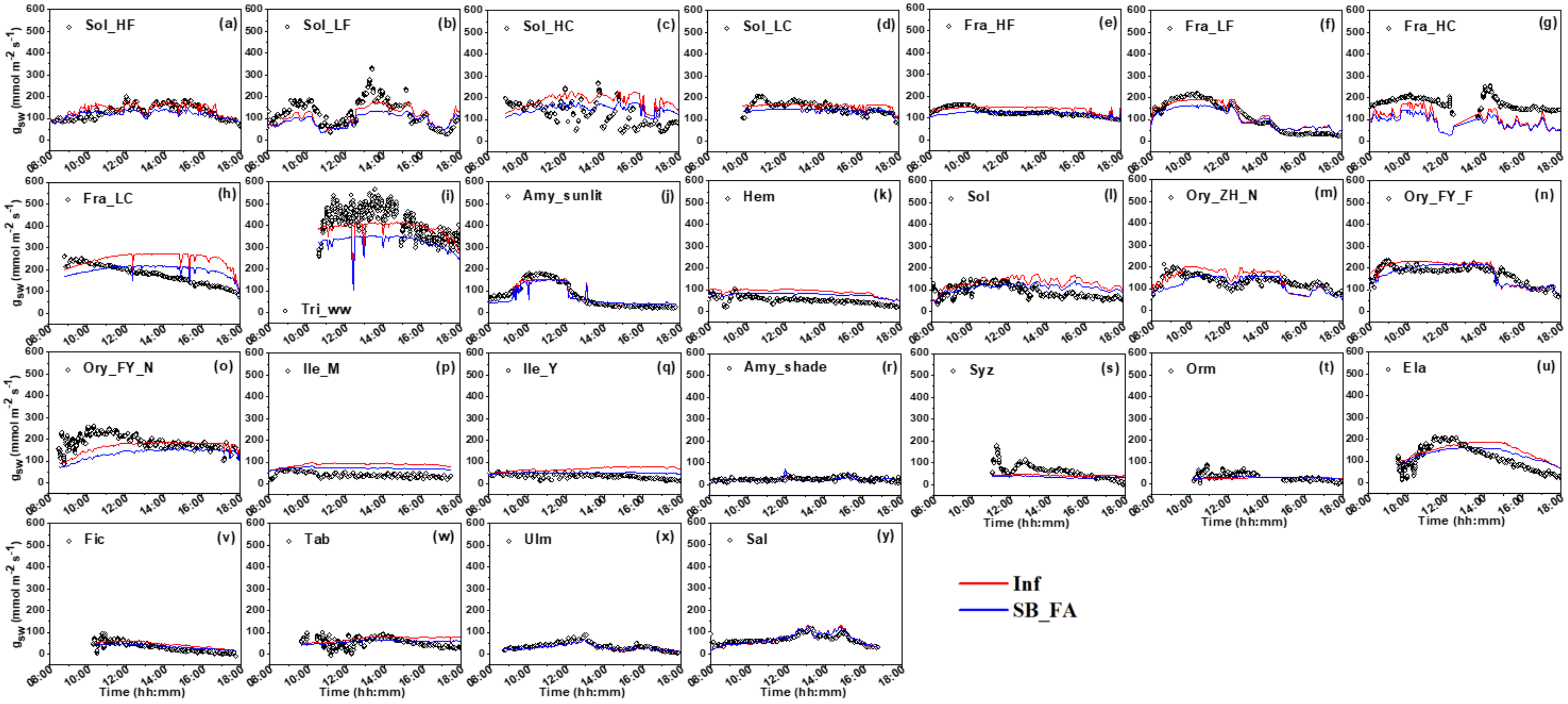

**Fig. S9** Comparisons in diurnal $A_n$ at *PAR* greater than 1500 μmol $m^{-2}$ $s^{-1}$ and across a wide range of leaf temperatures between observations (black squares) and predictions using the $g_m$ infinite model (red squares) and the $g_m$ finite model that is forced by the parameterization scheme SB_FA (blue squares).

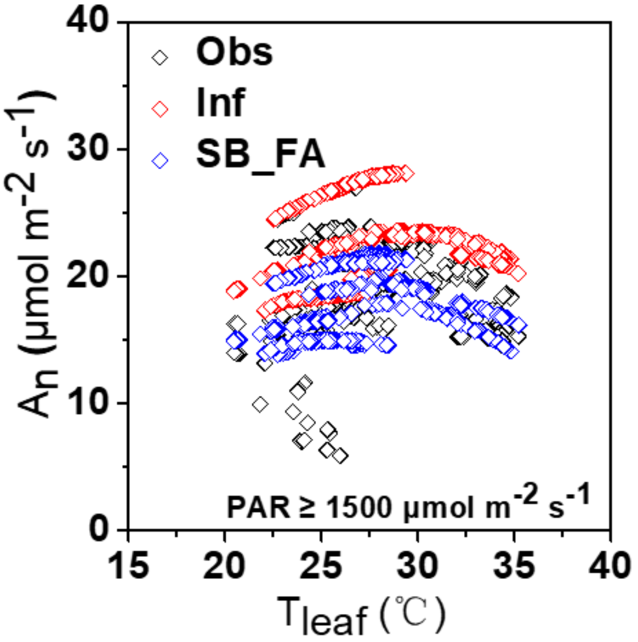

**Fig. S10** Statistical relationships for photosynthesis optimal temperature ($T_{optA}$) and the ratio of the maximum carboxylation rate to the maximum electron transport rate (*JV*r) (a), $T_{optA}$–$g_{m\_F,25}$ (b), and $T_{optA}$–activation term of of $g_{m\_F}$ ($\Delta H_a$) (c). The red solid line is the linear regression fit.

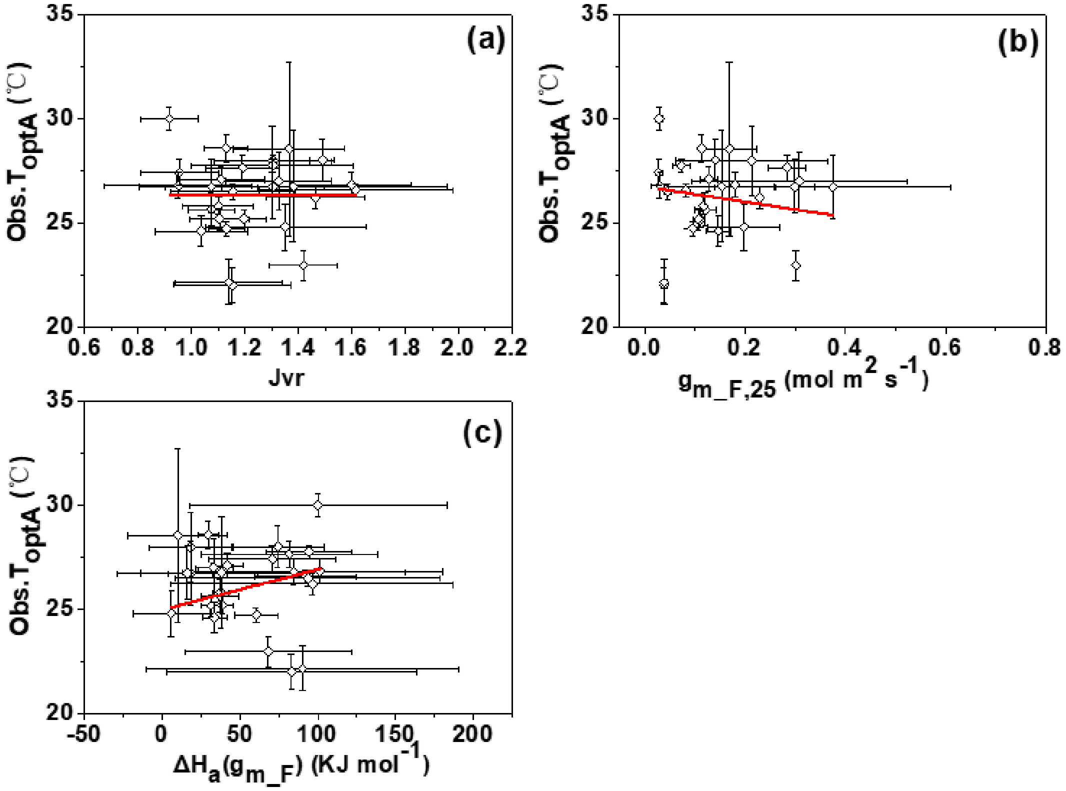

**Table S1** Temperature response characteristic parameters of $V_{cmax}$. Mean ± SD. TDF: tropical deciduous trees; EBF: evergreen broadleaf trees; TRF: tropical evergreen trees; DBF: deciduous broadleaf trees; $C_3$G: $C_3$ herbs and grasses; $C_3$C: $C_3$ crops; CV: coefficient of variation.

| PFT | Species | $V_{cmax\_B,25}$ (μmol $m^{-2}$ $s^{-1}$) | $V_{cmax\_S,25}$ (μmol $m^{-2}$ $s^{-1}$) | $V_{cmax\_SB,25}$ (μmol $m^{-2}$ $s^{-1}$) | $V_{a,25}$ (μmol $m^{-2}$ $s^{-1}$) | $\Delta H_a(V_{cmax\_B})$ (KJ $mol^{-1}$) | $\Delta H_a(V_{cmax\_S})$ (KJ $mol^{-1}$) | $\Delta H_a(V_{cmax\_SB})$ (KJ $mol^{-1}$) | $\Delta H_a(V_a)$ (KJ $mol^{-1}$) | $\Delta S(V_{cmax\_B})$ (J $K^{-1}$ $mol^{-1}$) | $\Delta S(V_{cmax\_S})$ (J $K^{-1}$ $mol^{-1}$) | $\Delta S(V_{cmax\_SB})$ (J $K^{-1}$ $mol^{-1}$) | $\Delta S(V_a)$ (J $K^{-1}$ $mol^{-1}$) |
|---|---|---|---|---|---|---|---|---|---|---|---|---|---|
| C3C | Sol_HF | 143.10±9.29 | 153.71±19.48 | 148.89±10.53 | 93.7±9.22 | 49.62±11.27 | 78.21±28.66 | 63.40±13.81 | 77.6±20.48 | 630.98±3.29 | 641.44±8.80 | 636.94±5.31 | 637.49±7.18 |
| | Sol_LF | 124.64±5.89 | 105.89±6.99 | 115.25±5.17 | 85.77±3.2 | 58.15±8.51 | 58.25±11.94 | 57.65±8.05 | 80.51±7.9 | 630.0±5.09 | 630.79±6.74 | 630.0±4.83 | 637.88±2.69 |
| | Sol_HC | 142.02±11.46 | 138.85±15.53 | 140.43±9.28 | 87.41±3.13 | 58.78±14.55 | 57.78±20.12 | 58.06±11.86 | 72.71±7.09 | 630.0±8.68 | 630.32±11.79 | 630.0±7.12 | 635.03±2.86 |
| | Sol_LC | 146.28±4.39 | 134.19±10.36 | 139.99±8.29 | 80.46±5.67 | 65.69±5.55 | 71.57±16.41 | 63.72±11.12 | 67.57±13.45 | 630.0±3.20 | 639.97±5.41 | 633.01±5.26 | 633.31±6.13 |
| | Tri_ww | 195.30±8.96 | 175.00±16.27 | 185.11±9.46 | 159.99±7.14 | 66.51±13.14 | 83.27±33.33 | 73.88±16.08 | 69.18±12.71 | 651.10±6.13 | 655.36±11.04 | 653.12±6.33 | 649.64±6.49 |
| | Gly_amb | 195.11±10.60 | 147.60±32.53 | 157.48±23.94 | 155.32±7.43 | 64.87±0.0 | 90.99±0.0 | 84.73±48.51 | 77.88±0 | 630.0±4.34 | 630.0±16.30 | 630.0±43.28 | 639.35±1.69 |
| **Mean** | | **157.74±29.99** | **142.54±22.94** | **147.86±23.07** | **110.44±36.85** | **60.6±6.46** | **73.35±13.47** | **66.91±10.51** | **74.24±5.22** | **633.68±8.54** | **637.98±9.92** | **635.51±9.05** | **638.78±5.74** |
| **CV(%)** | | **19.01** | **16.09** | **15.60** | **33.37** | **10.67** | **18.36** | **15.71** | **7.03** | **1.35** | **1.55** | **1.42** | **0.9** |
| C3G | Fra_HF | 193.16±7.65 | 208.86±14.43 | 200.64±8.88 | 108.37±13.14 | 72.68±7.92 | 88.15±18.37 | 79.80±9.89 | 82.64±24.59 | 635.95±3.05 | 645.01±5.11 | 640.71±3.07 | 634.76±9.66 |
| | Fra_LF | 194.18±16.40 | 192.82±21.14 | 193.48±12.85 | 104.31±9.23 | 56.87±15.12 | 58.95±20.12 | 57.91±12.04 | 76.02±17.88 | 630.13±9.02 | 632.59±10.02 | 631.41±6.52 | 635.8±6.83 |
| | Fra_HC | 203.92±5.28 | 195.65±15.07 | 200.13±7.69 | 99.9±8.56 | 66.63±4.81 | 83.24±16.14 | 72.57±7.43 | 79.56±16.66 | 630.0±2.76 | 636.89±5.68 | 632.76±3.43 | 631.28±8.25 |
| | Fra_LC | 209.52±7.45 | 233.46±15.59 | 220.88±9.42 | 109.54±3.05 | 65.02±6.99 | 92.53±21.64 | 77.25±10.03 | 78.78±6.05 | 636.90±2.67 | 649.05±5.97 | 643.19±3.03 | 639.5±1.96 |
| **Mean** | | **200.2±7.88** | **207.7±18.54** | **203.78±11.86** | **105.53±4.37** | **65.3±6.52** | **80.72±15.00** | **71.88±9.78** | **79.25±2.72** | **633.25±3.69** | **640.89±7.49** | **637.02±5.81** | **635.33±3.38** |
| **CV(%)** | | **3.94** | **8.93** | **5.82** | **4.14** | **9.98** | **18.58** | **13.61** | **3.43** | **0.58** | **1.17** | **0.91** | **0.53** |
| EBF | Ile_M | 48.15±7.66 | 43.76±6.46 | 46.05±4.85 | 21.42±0.82 | 75.93±28.29 | 81.50±26.95 | 78.52±18.99 | 75.77±8.62 | 630.0±15.96 | 630.0±14.80 | 630.0±10.58 | 647.01±2.61 |
| | Ile_Y | 50.33±3.06 | 46.77±3.38 | 48.72±2.39 | 17.87±1.14 | 58.59±9.59 | 71.36±12.53 | 64.87±8.14 | 78.12±14.36 | 630.0±6.00 | 630.0±7.24 | 630.0±4.89 | 645.85±4.28 |
| | E_cla | 135.57±11.21 | 129.04±10.87 | 134.23±10.32 | 88.98±13.6 | 88.33±15.72 | 78.52±19.15 | 78.18±14.94 | 80.31±29.58 | 630.0±12.35 | 638.01±8.28 | 630.0±12.36 | 630±24.19 |
| | E_dun | 101.19±7.72 | 78.21±15.43 | 90.78±8.99 | 60.69±9.12 | 62.47±15.27 | 133.29±216.47 | 82.48±25.38 | 73.26±29.59 | 630.0±14.09 | 653.56±49.25 | 641.16±9.10 | 630±25.57 |
| | E_ter | 147.63±14.48 | 117.06±22.59 | 132.33±13.41 | 91.88±14.06 | 67.82±19.30 | 75.51±37.55 | 71.32±19.85 | 68.95±30.06 | 630.0±17.16 | 630.0±31.88 | 630.0±17.28 | 630±26.54 |

| | | | | | | | | | | | | | |
|---|---|---|---|---|---|---|---|---|---|---|---|---|---|
| | E_sal | 140.78±15.76 | 93.93±9.75 | 117.40±10.29 | 87.51±19.33 | 101.96±59.68 | 122.44±68.41 | 110.15±50.57 | 98.18±123.55 | 650.92±15.15 | 651.27±16.54 | 650.94±12.60 | 653.46±29.78 |
| | E_mel | 172.76±11.02 | 163.39±22.64 | 169.87±12.11 | 129.21±19.09 | 70.69±12.61 | 86.74±31.12 | 72.92±14.01 | 76.84±39.45 | 630.0±11.06 | 638.11±12.85 | 630.0±12.02 | 642.05±13.66 |
| | E_cre | 156.16±14.70 | 132.01±18.72 | 144.96±12.32 | 94.78±16.13 | 65.71±18.11 | 120.41±97.06 | 80.88±20.22 | 85.3±31.84 | 630.0±16.50 | 651.79±22.81 | 640.28±7.69 | 630±25.66 |
| **Mean** | | **119.07±47.65** | **100.52±42.57** | **110.54±44.97** | **74.04±38.36** | **73.94±14.54** | **96.22±24.82** | **79.92±13.5** | **79.59±8.91** | **632.62±7.4** | **640.34±10.4** | **635.3±7.97** | **638.54±9.65** |
| **CV(%)** | | **40.02** | **42.35** | **40.69** | **51.81** | **19.66** | **25.80** | **16.89** | **11.19** | **1.17** | **1.62** | **1.25** | **1.51** |
| | Pop_CF | 60.75±13.40 | 47.46±4.62 | 53.90±6.16 | 67.64±2.8 | 86.65±49.92 | 100.17±22.75 | 93.25±26.32 | 97.43±14.66 | 630.0±54.45 | 630.0±23.19 | 630.0±27.74 | 651.81±3.87 |
| **DBF** | Pop_CF_N | 85.27±2.06 | 81.41±1.86 | 82.98±5.25 | 90.73±1.69 | 73.13±5.20 | 71.58±5.21 | 69.04±13.05 | 65.41±4 | 630.0±6.13 | 639.99±2.60 | 632.48±12.80 | 636.69±2.67 |
| | Pop_EO3 | 59.93±7.60 | 44.91±2.47 | 52.30±5.68 | 52.99±1.04 | 71.54±27.11 | 79.92±12.63 | 72.80±23.34 | 97.87±6.33 | 630.0±32.28 | 635.97±8.19 | 630.0±27.57 | 649.05±1.78 |
| | Pop_EO3_N | 82.50±7.60 | 77.78±5.81 | 79.33±6.07 | 91.1±6.91 | 66.03±19.18 | 95.36±24.01 | 72.12±17.28 | 98.53±26.71 | 630.0±23.64 | 649.39±6.75 | 638.52±9.56 | 651.39±7.09 |
| | Pop_CW | 136.08±50.84 | 115.10±10.31 | 137.69±22.19 | 146.33±50.23 | 142.08±0.00 | 100.41±0.00 | 84.25±42.50 | 59.84±0 | 652.08±6.13 | 630.05±4.97 | 630.0±34.88 | 630±21.25 |
| | Pop_WC | 180.02±37.79 | 172.92±41.60 | 187.52±26.67 | 136.92±45.47 | 100.0±0.00 | 110.0±0.00 | 96.09±41.29 | 56.17±0 | 633.04±8.99 | 634.40±9.07 | 630.0±32.87 | 630±20.86 |
| | Ulm | 112.02±8.34 | 110.57±8.00 | 111.32±5.75 | 105.82±11.53 | 81.98±13.57 | 82.20±13.70 | 81.92±9.57 | 76.36±19.04 | 632.51±5.38 | 634.24±4.97 | 633.31±3.64 | 630±8.96 |
| | Sal | 114.73±13.76 | 126.54±38.23 | 120.52±19.34 | 117.91±14.52 | 91.35±22.78 | 89.45±60.46 | 90.61±31.38 | 73.45±31.58 | 636.16±7.02 | 637.33±17.98 | 636.84±9.46 | 641.37±8.9 |
| **Mean** | | **103.91±40.73** | **97.09±42.98** | **103.2±45.77** | **101.18±32.24** | **89.1±24.15** | **91.14±12.72** | **82.51±10.37** | **78.13±17.65** | **634.22±7.54** | **636.42±6.24** | **632.64±3.39** | **640.04±9.74** |
| **CV(%)** | | **39.20** | **44.27** | **44.35** | **31.86** | **27.11** | **13.95** | **12.57** | **22.59** | **1.19** | **0.98** | **0.54** | **1.52** |
| **TRF** | Syz | 88.49±9.70 | 82.58±10.54 | 85.53±7.01 | 21.9±2.42 | 115.78±42.12 | 111.77±49.95 | 113.81±31.81 | 131.44±52.33 | 657.04±11.88 | 657.03±14.03 | 657.02±8.95 | 654.79±14.75 |
| | Orm | 92.48±6.76 | 71.55±11.09 | 82.31±6.69 | 15.39±2.26 | 55.02±15.87 | 29.97±24.89 | 43.22±15.08 | 84.76±36.8 | 643.31±5.57 | 632.80±12.94 | 639.37±5.86 | 643.61±10.4 |
| | Ela | 80.11±6.60 | 50.57±4.20 | 65.95±7.44 | 42.59±4.13 | 89.56±13.79 | 90.67±13.74 | 88.11±18.58 | 92.69±24.63 | 631.22±5.33 | 630.0±5.67 | 630.0±7.75 | 643.11±6.74 |
| | Fic | 55.38±8.64 | 42.70±13.39 | 49.32±8.63 | 29.62±3.87 | 119.42±90.10 | 121.27±182.73 | 123.14±101.27 | 139.35±66.86 | 658.82±21.53 | 655.0±43.97 | 657.76±24.23 | 661.74±16.15 |
| **Mean** | | **79.12±16.64** | **61.85±18.42** | **70.78±16.68** | **27.38±11.69** | **94.95±29.76** | **88.42±41.01** | **92.07±35.78** | **112.06±27.33** | **647.6±12.93** | **643.71±14.28** | **646.04±13.66** | **650.81±9.07** |
| **CV(%)** | | **21.04** | **29.78** | **23.56** | **42.7** | **31.34** | **46.38** | **38.86** | **24.39** | **2.00** | **2.22** | **2.11** | **1.39** |
| **TDF** | Tab | 82.99±15.03 | 71.59±14.31 | 77.23±10.07 | 24.73±5.33 | 136.13±85.69 | 143.86±83.53 | 140.04±57.80 | 119.52±69.44 | 655.38±22.50 | 655.91±22.21 | 655.67±15.27 | 645.6±17.39 |
| **Mean of all Species** | | **126.79±50.33** | **115.67±53.74** | **121.69±51.6** | **84.54±40.97** | **78.87±23.55** | **89.01±24** | **80.54±20.19** | **83.61±18.83** | **635.99±9.63** | **639.88±9.56** | **636.92±9.05** | **640.37±8.89** |

| | | | | | | | | | | | | |
|---|---|---|---|---|---|---|---|---|---|---|---|---|
| **CV(%) of all Species** | **39.69** | **46.46** | **42.40** | **48.46** | **29.86** | **26.97** | **25.07** | **22.52** | **1.51** | **1.49** | **1.42** | **1.39** |

**Table S2** Temperature response characteristic parameters of $J_{max}$. Mean ± SD. TDF: tropical deciduous trees; EBF: evergreen broadleaf trees; TRF: tropical evergreen trees; DBF: deciduous broadleaf trees; $C_3$G: $C_3$ herbs and grasses; $C_3$C: $C_3$ crops; CV: coefficient of variation.

| PFT | Species | $J_{max\_B,25}$ (μmol $m^{-2}$ $s^{-1}$) | $J_{max\_S,25}$ (μmol $m^{-2}$ $s^{-1}$) | $J_{max\_SB,25}$ (μmol $m^{-2}$ $s^{-1}$) | $J_{a,25}$ (μmol $m^{-2}$ $s^{-1}$) | $\Delta H_a(J_{max\_B})$ (KJ $mol^{-1}$) | $\Delta H_a(J_{max\_S})$ (KJ $mol^{-1}$) | $\Delta H_a(J_{max\_SB})$ (KJ $mol^{-1}$) | $\Delta H_a(J_a)$ (KJ $mol^{-1}$) | $\Delta S(J_{max\_B})$ (J $K^{-1}$ $mol^{-1}$) | $\Delta S(J_{max\_S})$ (J $K^{-1}$ $mol^{-1}$) | $\Delta S(J_{max\_SB})$ (J $K^{-1}$ $mol^{-1}$) | $\Delta S(J_a)$ (J $K^{-1}$ $mol^{-1}$) |
|---|---|---|---|---|---|---|---|---|---|---|---|---|---|
| | Sol_HF | 169.39±7.64 | 163.13±9.19 | 166.23±5.81 | 163.5±8.79 | 31.82±7.52 | 35.12±9.92 | 33.38±5.98 | 38.34±9.66 | 640.22±3.29 | 642.47±4.05 | 641.32±2.52 | 642.32±3.83 |
| $C_3$C | Sol_LF | 151.80±4.31 | 133.08±5.22 | 142.41±4.01 | 136.86±4.86 | 19.78±4.12 | 18.94±5.71 | 19.35±4.09 | 23.93±5.33 | 631.44±2.93 | 634.18±3.51 | 632.71±2.71 | 633.34±3.26 |
| | Sol_HC | 169.01±6.56 | 147.70±6.12 | 158.36±4.90 | 154.08±5.66 | 30.43±6.19 | 27.51±6.47 | 29.06±4.89 | 33.82±6.08 | 636.31±3.09 | 636.21±3.33 | 636.25±2.48 | 638.03±2.76 |
| | Sol_LC | 166.57±4.17 | 144.86±5.63 | 155.72±4.13 | 149.1±4.39 | 32.75±4.03 | 30.38±6.15 | 31.64±4.23 | 34.04±4.95 | 635.21±2.07 | 635.11±1.11 | 635.16±2.20 | 639.57±2.15 |
| | Tri_ww | 272.18±11.64 | 249.39±14.21 | 260.78±9.23 | 310.28±22.46 | 41.95±11.69 | 43.36±16.78 | 42.58±10.01 | 58.26±27.48 | 654.78±4.96 | 656.43±6.46 | 655.57±4.04 | 659.72±8.46 |
| | Gly_amb | 246.00±9.85 | 210.07±6.74 | 228.52±15.87 | 286.64±2.07 | 29.67±0.00 | 31.53±0.00 | 31.79±21.72 | 49.02±0 | 630.0±3.85 | 630.0±3.05 | 630.0±21.83 | 635.18±0.39 |
| **Mean** | | **195.83±50.12** | **174.71±45.42** | **185.34±47.68** | **200.07±77.05** | **31.07±7.09** | **31.14±8.1** | **31.3±7.48** | **39.57±12.23** | **637.99±8.99** | **639.07±9.41** | **638.5±9.18** | **641.36±9.54** |
| **CV(%)** | | **25.59** | **26.00** | **25.73** | **38.51** | **22.83** | **26.01** | **23.90** | **30.91** | **1.41** | **1.47** | **1.44** | **1.49** |
| | Fra_HF | 237.26±8.25 | 209.71±5.05 | 223.50±5.74 | 223.39±10.75 | 48.01±6.58 | 43.24±4.39 | 45.72±4.77 | 51.72±9.48 | 641.17±2.44 | 640.84±1.71 | 640.99±1.81 | 642.14±3.36 |
| $C_3$G | Fra_LF | 226.78±11.43 | 194.49±10.23 | 210.65±8.06 | 202.07±9.78 | 31.71±8.18 | 27.00±8.19 | 29.50±6.09 | 31.62±7.86 | 637.41±3.87 | 636.06±4.26 | 636.79±3.01 | 637.65±3.69 |
| | Fra_HC | 238.54±8.78 | 210.91±8.00 | 224.72±7.05 | 216.32±11.91 | 39.73±6.15 | 37.07±6.30 | 38.45±5.22 | 42.41±9.37 | 633.96±3.19 | 635.59±3.07 | 634.69±2.63 | 634.3±4.68 |
| | Fra_LC | 260.91±7.74 | 224.65±6.39 | 241.60±6.15 | 235.5±7.52 | 41.63±5.45 | 41.53±5.24 | 42.03±4.70 | 45.19±6.17 | 642.25±2.10 | 643.27±2.05 | 642.76±1.82 | 643.65±2.27 |
| **Mean** | | **240.87±14.36** | **209.94±12.33** | **225.12±12.7** | **219.32±13.96** | **40.27±6.72** | **37.21±7.29** | **38.93±6.95** | **42.73±8.38** | **638.7±3.78** | **638.94±3.74** | **638.81±3.72** | **639.43±4.27** |
| **CV(%)** | | **5.96** | **5.87** | **5.64** | **6.37** | **16.68** | **19.58** | **17.85** | **19.61** | **0.59** | **0.58** | **0.58** | **0.67** |
| | Ile_M | 48.64±3.82 | 58.07±3.54 | 53.33±2.69 | 46.22±1.78 | 31.53±9.45 | 32.74±7.44 | 32.18±6.12 | 37.99±5.4 | 630.0±7.55 | 630.0±5.87 | 630.0±4.85 | 644.94±2.41 |
| | Ile_Y | 48.77±1.53 | 57.66±2.48 | 53.10±1.90 | 41.52±1.29 | 29.99±3.76 | 27.95±4.98 | 28.53±4.18 | 35.99±4.17 | 636.57±2.13 | 630.0±4.14 | 632.99±2.86 | 643.55±1.92 |
| EBF | E_cla | 164.73±10.95 | 174.12±6.19 | 170.50±7.11 | 169.83±13.07 | 54.28±13.31 | 34.73±7.03 | 43.40±8.28 | 48.11±15.37 | 630.0±17.70 | 632.0±6.44 | 630.0±8.53 | 630±15.31 |
| | E_dun | 110.78±7.29 | 112.72±7.21 | 111.57±4.95 | 105.46±8.19 | 25.67±12.37 | 28.41±12.53 | 25.94±8.37 | 27.31±14.72 | 630.0±15.16 | 633.58±10.83 | 630.39±9.86 | 630±17.77 |
| | E_ter | 175.59±13.84 | 170.50±13.56 | 173.04±9.42 | 170.24±19.28 | 31.20±14.79 | 28.47±14.74 | 29.87±10.15 | 37.49±21.77 | 630.0±17.26 | 630.0±17.62 | 630.0±11.99 | 630±24.08 |
| | E_sal | 160.20±11.26 | 147.59±9.84 | 153.93±7.27 | 153.23±13.88 | 60.37±22.45 | 52.82±18.17 | 56.91±14.05 | 60.76±29.61 | 649.79±6.83 | 646.02±6.71 | 648.15±4.63 | 650.94±8.64 |
| | E_mel | 220.25±16. | 211.20±10. | 216.03±9.8 | 230.5±18.5 | 41.82±15.0 | 33.61±9.32 | 37.45±8.95 | 42.18±15.9 | 630.0±16.0 | 630.0±10.3 | 630.0±9.77 | 630±16.5 |

| | | | | | | | | | | | | | |
|---|---|---|---|---|---|---|---|---|---|---|---|---|---|
| | | 66 | 15 | 4 | 2 | 4 | | | 2 | 8 | 7 | | 6 |
| | E_cre | 185.95±13.71 | 183.36±11.11 | 184.13±8.73 | 184.04±18.17 | 33.77±13.54 | 35.49±12.33 | 32.78±8.90 | 40.74±18.58 | 630.0±16.01 | 638.86±6.98 | 633.34±7.85 | 630±20.62 |
| **Mean** | | **139.36±63.65** | **139.4±57.74** | **139.45±60.73** | **137.63±67.35** | **38.58±12.53** | **34.28±8.08** | **35.88±10.1** | **41.32±9.82** | **633.3±7.05** | **633.81±5.81** | **633.11±6.24** | **636.18±8.78** |
| **CV(%)** | | **45.67** | **41.42** | **43.55** | **48.94** | **32.49** | **23.56** | **28.14** | **23.77** | **1.11** | **0.92** | **0.99** | **1.38** |
| | Pop_CF | 89.01±4.70 | 82.80±3.31 | 85.95±2.70 | 91.86±1.58 | 28.09±8.64 | 34.47±6.83 | 31.14±5.26 | 37.27±3.1 | 630.0±14.02 | 630.0±10.55 | 630.0±8.33 | 638.65±2.16 |
| **DBF** | Pop_CF_N | 123.80±8.86 | 111.45±2.12 | 117.51±4.25 | 122.05±2.01 | 44.16±14.24 | 38.10±3.48 | 41.04±6.87 | 41.15±3.05 | 644.58±6.45 | 640.08±2.16 | 642.47±3.58 | 639.2±1.98 |
| | Pop_EO3 | 82.01±1.59 | 78.05±2.37 | 80.01±1.44 | 83.77±2.37 | 35.61±3.55 | 38.13±5.61 | 36.77±3.30 | 47.47±5.89 | 642.18±1.96 | 641.71±3.13 | 641.91±1.84 | 646.1±2.42 |
| | Pop_EO3_N | 114.41±4.71 | 106.79±4.99 | 110.60±3.46 | 118.55±6.14 | 31.53±7.11 | 34.98±8.37 | 33.18±5.50 | 42.53±10 | 638.40±5.26 | 640.55±5.16 | 639.48±3.71 | 642.9±5.03 |
| | Pop_CW | 192.10±15.45 | 176.24±11.28 | 185.54±11.74 | 242.7±55.26 | 28.00±0.00 | 28.10±0.00 | 28.53±10.34 | 30.89±0 | 630.0±6.00 | 630.0±4.78 | 630.0±10.89 | 630±17.96 |
| | Pop_WC | 212.94±57.29 | 206.75±55.53 | 226.4±27.28 | 319.67±120.29 | 67.45±0.00 | 76.81±0.00 | 59.27±25.55 | 38.76±0 | 632.34±13.17 | 637.13±8.83 | 630.0±22.93 | 630±25.93 |
| | Ulm | 149.45±6.34 | 131.25±3.92 | 140.34±4.32 | 140.03±5.69 | 52.40±8.48 | 57.21±6.46 | 54.66±6.37 | 66.19±9.06 | 637.44±3.13 | 640.19±2.17 | 638.77±2.24 | 640.59±2.87 |
| | Sal | 155.52±9.18 | 120.37±11.81 | 137.97±7.93 | 124.76±11.9 | 45.01±13.64 | 58.71±22.67 | 51.07±13.24 | 66.59±26.94 | 637.42±4.71 | 638.54±7.16 | 637.86±4.39 | 642.52±7.65 |
| **Mean** | | **139.91±46.67** | **126.71±44.47** | **135.54±49.68** | **155.42±82.31** | **41.53±13.67** | **45.81±16.63** | **41.96±11.63** | **46.36±13.23** | **636.55±5.41** | **637.28±4.69** | **636.31±5.44** | **638.74±5.88** |
| **CV(%)** | | **33.36** | **35.10** | **36.65** | **52.96** | **32.91** | **36.29** | **27.71** | **28.54** | **0.85** | **0.74** | **0.86** | **0.92** |
| **TRF** | Syz | 75.36±5.89 | 85.64±5.72 | 80.46±4.13 | 53.68±3.66 | 89.27±31.78 | 85.44±26.94 | 87.19±20.80 | 72.72±24.12 | 656.55±8.91 | 656.63±7.57 | 656.55±5.84 | 651.63±7.39 |
| | Orm | 88.92±4.30 | 101.95±8.08 | 95.51±4.87 | 52.79±3.42 | 114.61±23.39 | 139.68±18.47 | 135.05±14.02 | 35.11±12.66 | 670.79±5.29 | 679.13±4.83 | 676.86±3.46 | 644.18±5.17 |
| | Ela | 89.44±2.69 | 86.57±2.68 | 88.02±1.97 | 83.55±2.67 | 60.57±5.92 | 47.23±5.31 | 53.72±4.07 | 59.65±6.92 | 638.10±2.00 | 631.78±2.43 | 635.16±1.55 | 640.86±2.25 |
| | Fic | 56.09±5.13 | 56.62±7.25 | 56.52±4.49 | 51.51±4.03 | 94.96±57.00 | 80.97±59.71 | 89.66±44.03 | 94.49±49.68 | 660.24±13.58 | 656.18±15.11 | 658.63±10.70 | 660.5±11.81 |
| **Mean** | | **77.45±15.66** | **82.7±18.92** | **80.13±16.9** | **60.38±15.47** | **89.85±22.34** | **88.31±38.25** | **91.41±33.4** | **65.5±24.84** | **656.42±13.62** | **655.93±19.34** | **656.8±17.07** | **649.3±8.72** |
| **CV(%)** | | **20.22** | **22.88** | **21.09** | **25.62** | **24.86** | **43.31** | **36.54** | **37.92** | **2.08** | **2.95** | **2.60** | **1.34** |
| **TDF** | Tab | 71.55±10.51 | 70.76±10.88 | 71.16±7.25 | 57.21±9.7 | 117.65±67.58 | 117.22±71.76 | 117.45±47.20 | 120.64±72.97 | 657.24±17.39 | 657.03±18.47 | 657.14±12.15 | 658.09±18.8 |
| **Mean of all Species** | | **153.35±66.26** | **142.53±56.26** | **148.52±61.67** | **152.29±78.84** | **47.27±25.26** | **46.67±27.22** | **46.75±26.59** | **48.14±20.1** | **639.5±10.66** | **639.99±11.07** | **639.55±11.17** | **640.66±8.79** |
| **CV(%) of all Species** | | **43.21** | **39.47** | **41.52** | **51.77** | **53.43** | **58.31** | **56.88** | **41.75** | **1.67** | **1.73** | **1.75** | **1.37** |

**Table S3** Temperature response characteristic parameters of $g_m$. Mean ± SD. TDF: tropical deciduous trees; EBF: evergreen broadleaf trees; TRF: tropical evergreen trees; DBF: deciduous broadleaf trees; $C_3$G: $C_3$ herbs and grasses; $C_3$C: $C_3$ crops; CV: coefficient of variation.

| PFT | Species | $g_{m\_B,25}$ (mol $m^{-2}$ $s^{-1}$) | $g_{m\_S,25}$ (mol $m^{-2}$ $s^{-1}$) | $g_{m\_FA,25}$ (mol $m^{-2}$ $s^{-1}$) | $\Delta H_a(g_{m\_B})$ (KJ $mol^{-1}$) | $\Delta H_a(g_{m\_S})$ (KJ $mol^{-1}$) | $\Delta H_a(g_{m\_FA})$ (KJ $mol^{-1}$) | $\Delta S(g_{m\_B})$ (J $K^{-1}$ $mol^{-1}$) | $\Delta S(g_{m\_S})$ (J $K^{-1}$ $mol^{-1}$) | $\Delta S(g_{m\_FA})$ (J $K^{-1}$ $mol^{-1}$) |
|---|---|---|---|---|---|---|---|---|---|---|
| | Sol_HF | 0.26±0.05 | 0.35±0.10 | 0.11±0.01 | 40.87±35.71 | 123.32±73.48 | 51.5±22.82 | 641.91±13.86 | 684.19±17.13 | 654.07±7.55 |
| | Sol_LF | 0.32±0.04 | 0.32±0.08 | 0.11±0.00 | 90.83±56.82 | 11.84±35.55 | 55.68±15.1 | 658.72±15.89 | 643.6±16.4 | 655.33±4.78 |
| **C3C** | Sol_HC | 0.24±0.03 | 0.2±0.04 | 0.1±0.00 | 61.02±32.81 | 61.85±57.53 | 37.63±10.01 | 652.99±10.59 | 652.98±18.48 | 647.9±3.85 |
| | Sol_LC | 0.18±0.02 | 0.16±0.02 | 0.09±0.00 | 71.43±45.96 | 38.25±29.75 | 75.74±20.87 | 661.52±12.20 | 653.98±10.94 | 664.07±5.18 |
| | Tri_ww | 0.54±0.07 | 0.71±0.09 | 0.29±0.03 | 25.49±23.7 | 159.74±43.39 | 68.26±53.76 | 630±113.66 | 689.97±11.07 | 665.31±12.46 |
| | Gly_amb | 0.44±0.03 | - | 0.25±0.00 | 97.93±0.00 | - | 121.63±0.00 | 664.04±2.00 | - | 658.92±0.05 |
| **Mean** | | **0.33±0.14** | **0.35±0.22** | **0.16±0.09** | **64.59±28.11** | **79±61.13** | **68.41±29.26** | **651.53±13.16** | **664.94±20.71** | **657.6±6.55** |
| **CV(%)** | | **41.22** | **62.15** | **54.69** | **43.52** | **77.38** | **42.78** | **2.02** | **3.11** | **1.00** |
| | Fra_HF | 0.3±0.05 | 0.25±0.04 | 0.13±0.01 | 63.68±41.53 | 37.64±27.63 | 42.14±9.60 | 649.1±13.13 | 641.93±11.08 | 644.1±3.54 |
| **C3G** | Fra_LF | 0.22±0.02 | 0.22±0.03 | 0.14±0.01 | 44.04±16.22 | 27.59±23.61 | 49.95±19.92 | 646.85±5.92 | 641.27±10.46 | 653.17±6.74 |
| | Fra_HC | 0.22±0.01 | 0.22±0.01 | 0.12±0.00 | 45.76±11.36 | 25.21±9.42 | 29.34±5.98 | 642.47±4.21 | 630±7.05 | 637.98±2.78 |
| | Fra_LC | 0.22±0.01 | 0.18±0.01 | 0.12±0.00 | 37.47±10.25 | 17.34±5.6 | 37.24±4.90 | 645.27±4.03 | 637.86±3.32 | 647.64±1.96 |
| **Mean** | | **0.24±0.04** | **0.22±0.03** | **0.12±0.01** | **47.74±11.21** | **26.95±8.37** | **39.67±8.65** | **645.92±2.79** | **637.77±5.48** | **645.72±6.37** |
| **CV(%)** | | **16.51** | **14.59** | **8.23** | **23.49** | **31.05** | **21.80** | **0.43** | **0.86** | **0.99** |
| | Ile_M | 0.04±0.01 | 0.04±0.00 | 0.03±0.00 | 64.01±66.67 | 54.73±19.04 | 83.11±80.22 | 654.62±22.64 | 656.75±6.79 | 668.81±19.18 |
| | Ile_Y | 0.02±0.00 | 0.03±0.00 | 0.03±0.00 | 37.47±13.26 | 40.83±9.75 | 90.08±100.44 | 652.67±5.92 | 651.24±4.25 | 669.5±23.58 |
| | E_cla | 0.3±0.05 | 0.11±0.19 | 0.29±0.07 | 130.9±72.81 | 184.7±1534.23 | 147.28±68.53 | 675.33±21.74 | 652.25±337.35 | 679.07±24.31 |
| **EBF** | E_dun | 0.18±0.04 | 0.49±0.20 | 0.13±0.03 | 76.43±113.65 | 138.69±110.17 | 13.63±45.45 | 659.45±25.94 | 678.01±30.66 | 630±52.03 |
| | E_ter | 0.27±0.06 | 0.2±0.14 | 0.16±0.02 | 132.82±120.81 | 74.87±115.89 | 2.06±23.34 | 672.81±34.47 | 630±100.64 | 643.78±11.94 |
| | E_sal | 0.32±0.10 | 0.33±0.07 | 0.22±0.03 | 103.56±231.18 | 70.61±230.53 | 0.00±0.00 | 664.18±51.53 | 667.5±55.98 | 648.13±8.56 |
| | E_mel | 0.44±0.07 | 0.3±0.11 | 0.36±0.07 | 118.82±126.2 | 54.75±66.34 | 0.00±0.00 | 671.16±33.33 | 630±65.27 | 640.7±8.58 |
| | E_cre | 0.36±0.08 | 0.29±0.08 | 0.3±0.06 | 167.42±52.12 | 0.00±0.00 | 61.24±75.83 | 680.16±19.63 | 630±52.05 | 655.27±19.22 |
| **Mean** | | **0.24±0.15** | **0.23±0.16** | **0.19±0.13** | **103.93±42.39** | **77.4±58.16** | **49.68±54.66** | **666.3±10.09** | **649.47±18.29** | **654.41±16.82** |
| **CV(%)** | | **62.05** | **71.03** | **66.06** | **40.79** | **75.14** | **110.03** | **1.52** | **2.82** | **2.57** |
| | Pop_CF | 0.71±0.48 | - | 0.19±0.02 | 105.4±435.87 | - | 101.59±78.85 | 662.91±101.71 | - | 663.03±18.44 |
| | Pop_CF_N | 0.71±0.10 | - | 0.25±0.04 | 121.84±41.89 | - | 96.37±90.41 | 680.77±9.54 | - | 661.97±21.45 |
| **DBF** | Pop_EO3 | 0.24±0.07 | - | 0.08±0.01 | 98.23±213.3 | - | 92.19±32.53 | 664.33±49.53 | - | 662.37±7.74 |
| | Pop_EO3_N | 0.76±0.15 | - | 0.16±0.02 | 106.47±173.41 | - | 74.18±29.86 | 666.39±39.60 | - | 651.71±8.74 |
| | Pop_CW | 0.29±0.00 | 0.52±0.28 | 0.38±0.00 | 14.82±0.00 | 0.00±0.00 | 63.41±0.00 | 641.73±0.60 | 679.03±8.18 | 638.04±0.04 |
| | Pop_WC | 0.17±0.01 | 0.19±0.00 | 0.34±0.03 | 0.00±0.00 | 7.48±0.00 | 21.75±0.00 | 630±7.15 | 633.41±1.33 | 638.26±4.77 |
| | Ulm | - | - | 0.28±0.04 | - | - | 81.58±56.72 | - | - | 654.40±16.20 |
| | Sal | - | - | 0.30±0.04 | - | - | 16.02±45.11 | - | - | 644.02±14.11 |
| **Mean** | | **0.48±0.27** | **0.35±0** | **0.25±0.1** | **74.46±52.71** | **3.74±0** | **68.39±32.95** | **657.69±18.45** | **656.22±0** | **651.72±10.56** |
| **CV(%)** | | **56.70** | **66.59** | **40.25** | **70.79** | **141.42** | **48.19** | **2.80** | **4.92** | **1.62** |
| | Syz | 0.03±0.01 | 0.04±0.01 | 0.03±0.00 | 54.92±61.27 | 54.38±43.49 | 70.88±40.84 | 630±30.14 | 630±21.73 | 659.28±10.93 |
| **TRF** | Orm | 0.02±0.00 | 0.03±0.00 | 0.03±0.00 | 33.78±15.97 | 52.32±21.87 | 84.47±71.77 | 637.99±7.09 | 640.87±21.57 | 666.11±16.64 |
| | Ela | 0.1±0.01 | 0.28±0.18 | 0.06±0.00 | 52.09±17.38 | 85.17±193.52 | 94.4±27.56 | 653.61±5.54 | 656.99±56.85 | 662.48±6.79 |
| | Fic | 0.06±0.01 | 0.08±0.01 | 0.03±0.00 | 115.89±61.04 | 115.14±52.61 | 93.65±85.04 | 668.19±14.04 | 667.75±12.17 | 664.76±19.37 |
| **Mean** | | **0.05±0.04** | **0.11±0.11** | **0.04±0.02** | **64.17±35.73** | **76.75±29.68** | **85.85±10.95** | **647.45±16.95** | **648.9±16.76** | **663.16±2.99** |
| **CV(%)** | | **71.51** | **107.63** | **40.00** | **55.68** | **38.67** | **12.76** | **2.62** | **2.58** | **0.45** |

| | | | | | | | | | | |
|---|---|---|---|---|---|---|---|---|---|---|
| **TDF** | Tab | 0.03±0.01 | 0.03±0.03 | 0.02±0.00 | 114.17±84.52 | 95.72±169.64 | 100.06±79.29 | 658.37±21.51 | 634.44±58.56 | 659.01±19.5 |
| **Mean of all Species** | | **0.28±0.2** | **0.23±0.17** | **0.17±0.11** | **76.81±40.91** | **63.84±50.42** | **63.13±36.9** | **655.78±14.46** | **650.58±18.72** | **654.49±11.36** |
| **CV(%) of all Species** | | **74.15** | **73.26** | **67.30** | **53.26** | **78.98** | **58.45** | **2.20** | **2.88** | **1.74** |

**Methods S1 The explicit clarity on the parameter values assumed for each parameter estimation method**

**(1) Input parameters for the $g_m$ infinite model**

**Table S4** Input parameters of the $g_m$ infinite photosynthesis-transpiration coupled model. Fixed values of model input parameters were referred to Harley and Tenhunen (1991), Xue *et al*. (2017), Knauer et al. (2019) and Kumarathunge *et al*. (2019).

| Input parameters | Value | Unit | Parameter estimation and expression |
|---|---|---|---|
| $V_{a,25}$ | - | μmol $m^{-2}$ $s^{-1}$ | $A_n$/$C_i$ curves, Eqn. 7 |
| $\Delta H_a$ ($V_a$) | - | J $mol^{-1}$ | |
| $\Delta S$ ($V_a$) | - | J $K^{-1}$ $mol^{-1}$ | |
| $\Delta H_d$ ($V_a$) | 200000 | J $mol^{-1}$ | Fixed value |
| $J_{a,25}$ | - | μmol $m^{-2}$ $s^{-1}$ | $A_n$/$C_i$ curves, Eqn. 7 |
| $\Delta H_a$ ($J_a$) | - | J $mol^{-1}$ | |
| $\Delta S$ ($J_a$) | - | J $K^{-1}$ $mol^{-1}$ | |
| $\Delta H_d$ ($J_a$) | 200000 | J $mol^{-1}$ | Fixed value |
| $g_{fac}$ | - | | Diurnal gas exchange, Eqn. 8 |
| $g_{min}$ | - | mmol $m^{-2}$ $s^{-1}$ | |
| $g_{max}$ | 900 | mmol $m^{-2}$ $s^{-1}$ | Fixed value |
| $O_2$ | 210 | mmol $mol^{-1}$ | Fixed value |
| $\alpha$ | 0.24 | unitless | Fixed value |
| $\Theta$ | 0.85 | unitless | Fixed value |
| $R_{d25}$ | - | μmol $m^{-2}$ $s^{-1}$ | Gas exchange measurement |
| $\Delta H_a$ ($R_d$) | - | J $mol^{-1}$ | Xue *et al.* (2017) |
| $\Delta H_a$ ($K_o$) | 36380 | J $mol^{-1}$ | |
| $c$ ($K_o$) | 20.30 | mmol $mol^{-1}$ | |
| $\Delta H_a$ ($K_c$) | 79430 | J $mol^{-1}$ | Bernacchi *et al.* (2001) |
| $c$ ($K_c$) | 38.05 | μmol $mol^{-1}$ | |
| $\Delta H_a$ ($\Gamma$*) | 37830 | J $mol^{-1}$ | |
| $c$ ($\Gamma$*) | 19.02 | μmol $mol^{-1}$ | |

**(2) $g_m$ estimation using the chlorophyll fluorescence-gas exchange method**

Mesophyll conductance was estimated by using the variable $J$ method ($g_{m\_F}$) proposed by Harley *et al*. (1992). Assimilation rate and ETR in the region from 240 to 600 μmol $mol^{-1}$ were used, because (1) $g_m$ estimates over this range of intercellular $CO_2$ concentration ($C_i$) values are reliable, since photosynthesis is limited by regeneration of RuBP and limitations by triose phosphate utilization that usually occur only at higher $CO_2$ concentrations (Harley *et al*., 1992, Flexas *et al*., 2007, Sharkey *et al*., 2007); and (2) the determinations are appropriate for the values observed for $C_i$ under natural field conditions (ca. 250 μmol $mol^{-1}$). Photosynthetic values at $C_i$ of ca. 250 μmol $mol^{-1}$ were used for $g_m$ estimation, as follows:

$$g_{m\_F} = \frac{A_n}{C_i - \frac{\Gamma^*(J_F + 8(A + R_d))}{J_F - 4(A + R_d)}} \qquad (Eqn\ S1)$$

$$J_F = \Phi_{\text{PSII}} Q \alpha \beta \qquad \text{(Eqn S2)}$$

$$\Phi_{PS\,\text{II}} = \frac{F_m' - F_s}{F_m'} \qquad (Eqn\ S3)$$

where $J_F$ was the linear electron transport rate, $\alpha$ here denoted the light absorption capacity of the leaves (generally taken as 0.85), $\beta$ reflected the distribution ratio of activating light between light system I and II (PSI and PSII), and generally took a value of 0.5. $\Phi_{PSII}$ was the actual photochemical efficiency of photosystem II, which was calculated by the light adapted maximum ($F_m$') and steady-state ($F_s$) fluorescence yields of PSII.

**(3) $g_m$ estimation using the anatomy method**

The mesophyll conductance estimated by the anatomy method ($g_{m\_A}$) includes the gas phase conductance of $CO_2$ diffusion from the sub-stomata to the cell wall ($g_{ias}$) and the liquid phase conductance from the cell wall to the carboxylation site of $CO_2$ in the chloroplast ($g_{liq}$) (Evans *et al.*, 1994):

$$g_{m\_A} = \frac{1}{\frac{1}{g_{ias}} + \frac{RT_L}{H \cdot g_{liq}}} \qquad (Eqn\ S4)$$

where $H$ is the Henry coefficient (Pa $m^3$ $mol^{-1}$), [$H = 1/0.034 \times \text{EXP}(-2400 \times (1/\text{T}_L - 1/\text{T}_\text{n})) \times 101.325$] (Table S5), $T_L$ is the leaf temperature (K), and $T_n$ is the standard temperature (298.16K). [$H/(R \times T_L)$] is the dimensionless form of Henry's coefficient, which converts liquid phase conductance into equivalent gas phase conductance (Laisk *et al.*, 2002; Niinemets and Reichstein, 2003a; Tosens *et al.*, 2012). $g_{ias}$ and $g_{liq}$ were calculated from leaf structural data extracted from ultra-thin and semi-thin sections of leaves, referring to Tosens *et al.* (2012), Tomas *et al*. (2013), and Han *et al*. (2018).

The $g_{ias}$ could be expressed as follows:

$$g_{ias} = \frac{1}{r_{ias}} = \frac{D_a \cdot f_{ias}}{\Delta L_{ias} \cdot \varsigma} \qquad \text{(Eqn S5)}$$

Where $D_a$ is the $CO_2$ diffusion coefficient in the gas phase ($1.51 \times 10^{-5}$ $m^2$ $s^{-1}$ at 25°C), $\Delta L_{ias}$ is the diffusion length (m), generally being half the mesophyll thickness, $\varsigma$ is the diffusion tortuosity (m $m^{-1}$), which could be obtained from the leaf cross section (Niinemets and Reichstein, 2003a). $f_{ias}$ is the fraction of leaf air space, which could be expressed as follows:

$$f_{ias} = 1 - \frac{\Sigma S_s}{t \cdot w} \quad \text{(Eqn S6)}$$

where $\Sigma S_s$ is the total surface area of the mesophyll cells, $t$ is the thickness of the mesophyll (m), $w$ is the width of the section (m).

The $g_{\text{liq}}$ could be expressed as follows:

$$g_{liq} = \frac{1}{\left(\frac{1}{g_{cw}} + \frac{1}{g_{pl}} + \frac{1}{g_{ct}} + \frac{1}{g_{en}} + \frac{1}{g_{st}}\right)} \cdot \frac{S_c}{S} \quad \text{(Eqn S7)}$$

where $g_{cw}$, $g_{pl}$, $g_{ct}$, $g_{en}$ and $g_{st}$ are the partial conductance of the cell wall, plasmalemma, cytosol, chloroplast envelope, and chloroplast stroma, respectively; and $S_c/S$ was the exposed chloroplast to leaf area ratio that could be expressed as,

$$\frac{S_c}{S} = \frac{L_c}{L_m} \cdot \frac{S_m}{S} \quad \text{(Eqn S8)}$$

where $L_c$ is the total length of chloroplast surface area facing the intercellular air space in the tissue section (m); $L_m$ is the total length of the mesophyll cells facing the intercellular air space in the tissue section (m); $S_m/S$ is the exposed mesophyll cells to leaf area ratio, which can be obtained as follows:

$$\frac{S_m}{S} = \frac{L_m}{w}\gamma \quad \text{(Eqn S9)}$$

where γ is the curvature correction factor, which is calculated for each species by measuring the width and height of palisade and spongy cells and an average width/height ratio (Thain, 1983). $g_{cw}$, $g_{ct}$, and $g_{st}$ could be calculated by the following equation:

$$g_i = \frac{1}{r_i} = \frac{r_{f,i} \cdot D_w \cdot p_i}{\Delta L_i} \quad \text{(Eqn S10)}$$

where $D_w$ is the $CO_2$ diffusion coefficient in the liquid phase ($1.79 \times 10^{-9}$ $m^2$ $s^{-1}$ at 25°C); $\Delta L_i$ is the length of the diffusion path in the corresponding component of the diffusion pathway (m), including $L_{cw}$, $L_{ct}$, and half of $L_{st}$; $r_{f,i}$ is the reduction of $D_w$ compared with free diffusion in water; and $p_i$ is its effective porosity ($m^3$ $m^{-3}$). For the cell wall, $r_{f,i}$ is taken as1.0 and $p_i$ is taken as 0.3; for cytosol and stroma, $r_{f,i}$ is taken as 0.3 and $p_i$ was taken as 1.0. Both plasmalemma and chloroplast envelope had phospholipid bilayers, their conductance is estimated to be 0.0035 m $s^{-1}$ (Niinemets and Reichstein, 2003b; Rondeau-Mouro *et al.*, 2008; Tosens *et al.*, 2012; Tomas *et al.*, 2013). Finally, convert mesophyll conductance into mole units by using the following equation:

$$g_{ias}[molm^{-2}s^{-1}] = \frac{g_{ias}[ms^{-1}]44.6 \cdot 273.16P}{(273.16 + T_L)(101325)} \quad \text{(Eqn S11)}$$

where $P$ was the air pressure (Pa).

**Table S5** Parameters of the anatomy method used to estimate mesophyll conductance. Fixed values were referred to Syvertsen *et al*. (1995), Tosens *et al*. (2012), Tomas *et al*. (2013), and Xiong and Flexas (2021).

| Parameter | Trait | Value | Unit | Reference |
|---|---|---|---|---|
| $H$ | Henry coefficient | - | Pa $m^{-3}$ $mol^{-1}$ | Han *et al.* (2019) |
| $T_L$ | Leaf temperature | - | K | - |
| $D_a$ (25°C) | $CO_2$ diffusion coefficient in the gas phase | $1.51 \times 10^{-5}$ | $m^2$ $s^{-1}$ | Fixed value |
| $f_{ias}$ | Fraction of leaf air space | - | - | The semi-thin section |
| $\Delta L_{ias}$ | Half of mesophyll thickness | - | m | |
| $\varsigma$ | Diffusion tortuosity | 1.57 | m $m^{-1}$ | Fixed value |
| $\Sigma S_s$ | The total surface area of the mesophyll cells | - | $m^2$ | The semi-thin section |
| $t$ | Mesophyll thickness | - | m | |
| $w$ | Section width | - | m | |
| $L_m$ | The total length of the mesophyll cells facing the intercellular air space in the tissue section | - | m | |
| $L_c$ | The total length of chloroplast surface area facing the intercellular air space in the tissue section | - | m | The ultra-thin section |
| $\gamma$ | Curvature correction factor | - | - | (Thain, 1983) |
| $L_{cw}$ | Cell wall thickness | - | m | The ultra-thin section |
| $L_{ct}$ | Average distance between the chloroplast and the cell wall | - | m | |
| $L_{st}$ | Half of chloroplast thickness | - | m | |
| $S_c/S$ | The exposed chloroplast to leaf area ratio | - | - | The semi-thin section and the ultra-thin section |
| $S_m/S$ | The exposed mesophyll cells to leaf area ratio | - | - | The semi-thin section |
| $D_w$ (25°C) | $CO_2$ diffusion coefficient in the liquid phase | $1.79 \times 10^{-9}$ | $m^2$ $s^{-1}$ | Fixed value |
| $g_{cw}$ | The conductance of cell wall | - | m $s^{-1}$ | - |
| $g_{ct}$ | The conductance ofcytosol | - | m $s^{-1}$ | - |
| $g_{st}$ | The conductance ofchloroplast stroma | - | m $s^{-1}$ | - |
| $g_{pl}$ | The conductance ofplasmalemma | 0.0035 | m $s^{-1}$ | Fixed value |
| $g_{en}$ | The conductance ofchloroplast envelope | 0.0035 | m $s^{-1}$ | Fixed value |
| $r_{f,i}$ | The reduction of $D_w$ compared with free diffusion in water | 1.0 for cell wall; 0.3 for cytosol and stroma | - | Tosens *et al*. (2012) |
| $p_i$ | Effective porosity | 0.3 for cell wall; 1.0 for cytosol and stroma | - | |

**(4) The Rubisco kinetic parameters for the Bayesian retrieval algorithm and the Sharkey online calculator**

For the parameterization of the $g_m$ finite model, we used the Bayesian retrieval algorithm (Zhu *et al.*, 2011; Han *et al.*, 2020) and the Sharkey online calculator (Sharkey *et al.*, 2007) to estimate $V_{cmax}$, $J_{max}$, and $g_m$. The Farquhar, von Caemmerer & Berry (1980) (FvCB) model of $C_3$ photosynthesis was applied in both two methods. The algorithm theory of the Bayesian inversion method is based on the Farquhar photosynthesis model that determines $V_{cmax}$, $J_{max}$, and $g_m$ through a quadratic relationship for $A_n$ and $C_i$, where $C_i$ is a function of $C_c$, $A_n$, and $g_m$. It combines a priori probability density function (PDF) developed according to the possible ranges of $V_{cmax}$, $J_{max}$, and $g_m$ to generate a posterior PDF and provides optimized estimation values for $V_{cmax}$, $J_{max}$, and $g_m$ (Zhu *et al.*, 2011; Han *et al.*, 2020). The Sharkey online calculator uses least square regression to exclusively perform nonlinear fitting on the Rubisco carboxylation stage and the RuBP regeneration stage according to the $A_n/C_i$ curve and generates the required photosynthetic parameters (Sharkey *et al.*, 2007). The Rubisco kinetic parameters used by the Bayesian retrieval algorithm and the Sharkey online calculator to estimate photosynthetic parameters ($V_{cmax}$, $J_{max}$, and $g_m$) were shown in Table S6.

**Table S6** Datasets of $K_o$, $K_c$ and $\Gamma^*$ used by the Bayesian retrieval algorithm and the Sharkey online calculator to estimate photosynthetic parameters ($V_{cmax}$, $J_{max}$, and $g_m$). $\Delta H_a$ ($K_o$), $\Delta H_a$ ($K_c$) and $\Delta H_a$ ($\Gamma^*$): the activation energy for $K_o$, $K_c$, and $\Gamma^*$, respectively; $K_{o,25}$, $K_{c,25}$, and $\Gamma^*_{25}$: $K_o$, $K_c$, and $\Gamma^*$ values at 25°C, respectively.

| Parameter estimation method | Parameters | Value | Unit | Reference |
|---|---|---|---|---|
| the Bayesian retrieval algorithm | $\Delta H_a$ ($K_o$) | 35950 | J mol$^{-1}$ | Zhu *et al.* (2011) and |
| | $K_{o,25}$ | 16.58 | kPa | Han *et al.* (2020) |
| | $\Delta H_a$ ($K_c$) | 79970 | J mol$^{-1}$ | |
| | $K_{c,25}$ | 27.24 | Pa | |
| | $\Delta H_a$ ($\Gamma^*$) | 26800 | J mol$^{-1}$ | |
| | $\Gamma^*_{25}$ | 3.74 | Pa | |
| the Sharkey online calculator | $\Delta H_a$ ($K_o$) | 23720 | J mol$^{-1}$ | Sharkey *et al.* (2007) |
| | $K_{o,25}$ | 16.58 | kPa | |
| | $\Delta H_a$ ($K_c$) | 80990 | J mol$^{-1}$ | |
| | $K_{c,25}$ | 27.24 | Pa | |
| | $\Delta H_a$ ($\Gamma^*$) | 24460 | J mol$^{-1}$ | |
| | $\Gamma^*_{25}$ | 3.74 | Pa | |

**Methods S2** Plant materials used for photosynthesis and anatomical structure measurements were collected in Lanzhou of Gansu and Xishuangbanna of Yunnan Provinces, China. The Gansu province has a temperate continental climate, and the main forest type in this region is the temperate deciduous forest. The annual mean air temperature and the annual mean precipitation in this region are 6.6℃ and 300-400mm, respectively. Xishuangbanna is located in the northern tropical monsoon climate zone, with an annual mean air temperature of 21.5℃ and an annual mean precipitation of 1557 mm. Tropical rain forest is prevalent in this area. Experimental materials used were as follows: sunlit leaves of *Hemerocallis fulva* (abbreviated as Hem); sunlit and shaded leaves of the flowering plum *Amygdalus triloba* (abbreviated as Amy_sunlit and Amy_shade, respectively) obtained in July 2019 from the campus of Lanzhou University (36.08N, 103.98E); sunlit leaves of *Oryza sativa* cv. Zhenghan10 and *Oryza sativa* L. cv. Fangyuan grown under plastic film mulching and control conditions (abbreviated as Ory_ZH_F, Ory_ZH_N, Ory_FY_F, and Ory_FY_N, respectively); sunlit leaves of *Solanum tuberosum* (abbreviated as Sol) obtained in July and August 2019 from the Dingxi National Experimental Station of the China Meteorological Administration (35.933N, 105.0E); young and mature leaves of *Ilex chinensis* (abbreviated as Ile_Y and Ile_M, respectively) obtained in April 2020 from the campus of Lanzhou University; sunlit leaves of field-grown *Solanum tuberosum* and *Fragaria ananassa* under warming + fertilization conditions, no warming + fertilization conditions, warming + no fertilization conditions, and no warming + no fertilization conditions (abbreviated as Sol_HF, Fra_HF, Sol_LF, Fra_LF, Sol_HC, Fra_HC, Sol_LC, and Fra_LC, respectively) collected from July to September 2020 at Gansu Provincial Station for Dryland Agricultural Ecosystem Observation and Research (36.04N, 104.38E); sunlit leaves of five representative tropical tree species, namely *Syzygium polypetaloideum* Merrill & LM Perry (abbreviated as Syz), *Ormosia henryi* Prain (abbreviated as Orm), *Elaeocarpus hainanensis* Oliv (abbreviated as Ela), *Ficus tinctoria* Forst. F. subsp. Gibbose (BI.) Corner (abbreviated as Fic), and *Tabebuia heterophylla* (DC.) Britt (abbreviated as Tab), obtained in March and April 2021 from the Xishuangbanna Tropical Botanical Garden of the China Academy of Sciences (21.92N, 101.25E); and sunlit leaves of *Ulmus pumila* and *Salix matsudana* (abbreviated as Ulm and Sal, respectively) obtained in June 2021 from the Lanzhou University campus. The temperature-increasing treatment for *Solanum tuberosum* and *Fragaria ananassa* was performed using an open-top warming chamber (OTC), which showed an average monthly temperature increase of 4.1$^{o}$C and a relative humidity difference of less than 3.5% between the inside and outside of the OTC. The $A_n/C_i$ curve plus chlorophyll fluorescence data at different leaf temperatures and the diurnal gas exchange data were collected using those plant materials. Additionally, the following datasets regarding the $A_n/C_i$ curve and chlorophyll fluorescence data at different leaf temperatures were collected for analysis: *Glycine max* cv. NC-Roy (abbreviated as Gly_amb) from Xu *et al*. (2016); *Populus balsamifera* under two experimental conditions (abbreviated as Pop_CW and Pop_WC, respectively) from Silim *et al*. (2010); *Triticum asetivum* cv. Suan (abbreviated as Tri_ww) from Xue *et al*. (2016b); *Populous deltoides* cv. 55/56 and *P. deltoides* cv. Imperial under four experimental conditions (abbreviated as Pop_CF, Pop_CF_N, Pop_EO3, and Pop_EO3_N, respectively) from Xu *et al*. (2020); and six eucalyptus species, namely *Eucalyptus cladocalyx*, *Eucalyptus dunnii*, *Eucalyptus tereticornis*, *Eucalyptus saligna*, *Eucalyptus melliodora*, and *Eucalyptus crebra* (abbreviated as E_cla, E_dun, E_ter, E_sal, E_ta, and E_cre, respectively) from Kumarathunge *et al*. (2019).

**Methods S3** Measurements of the $A_n/C_i$ curve plus chlorophyll fluorescence at leaf temperatures ranging from 10-15 to 40°C were performed using a portable gas exchange and fluorescence system (GFS3000, Heinz Walz GmbH, Effeltrich, Germany). The relative humidity and light intensity inside the 2.0 cm × 4.0 cm leaf chamber were set at 50% and 1800 μmol $m^{-2}$ $s^{-1}$, respectively. The sampled leaves were acclimated to the microenvironment of the leaf chamber for 15–30 min (the $CO_2$ concentration at 400 ppm and the saturated light intensity at 1800 μmol $m^{-2}$ $s^{-1}$) before initiating the $CO_2$ concentration gradient of 1500, 1200, 900, 600, 400, 300, 200, 100, and 50 ppm. $A_n$ and stomatal conductance ($g_{sw}$) as well as chlorophyll fluorescence parameters were recorded when they became stable at each $CO_2$ concentration. The $A_n/C_i$ curve and simultaneous chlorophyll fluorescence measurements at each leaf temperature for each species and/or experimental treatment were repeated at least three times until the acquisition of high-quality data. Empty leaf chamber corrections before measurements were performed and other operational information were obtained according to the method reported by Xue *et al*. (2016). Two healthy leaves adjacent to the ones used for the $A_n/C_i$ curve and chlorophyll fluorescence measurements were chosen for the diurnal gas exchange measurement. The methods for the diurnal gas exchange measurement reported by Xue *et al*. (2017) were used in this study. The $A_n/C_i$ curve with chlorophyll fluorescence measurement at different leaf temperatures were assessed for plant species in 2020 and 2021, and the measurement at only 25°C was performed for the plant species in 2019.

**Methods S4** *In-situ* leaves were sampled to determine the leaf anatomical structure immediately after the gas exchange measurement. The pre-processing procedure for semi-thin and ultra-thin sections was as follows: firstly, a 4.0 mm × 1.2 mm segment without the veins was excised from at the middle portion of the leaves; the excised segment was placed in 2.5% (volume concentration) glutaraldehyde fixative solution (prepared using 0.1 M phosphate buffer, pH = 7.0–7.2) for fixation; a circulating water vacuum pump was then used to make the sample sink to the bottom; finally the process samples were stored in a refrigerator at 4°C. The pre-processed samples were sent to the Wuhan Saiweier Biotechnology Company for obtaining the semi-thin and ultra-thin sections. Leaf structural traits including the total length of the mesophyll cell wall facing the intercellular space ($l_s$), mesophyll thickness ($t$), intercellular space area, leaf cross-sectional area from the semi-thin section, the cell wall thickness ($T_{cw}$), chloroplast thickness ($L_{st}$), the average distance between the chloroplast and the cell wall ($L_{ct}$), the total length of the total mesophyll cells facing the intercellular space ($l_m$), and the total length of the chloroplast facing the intercellular space ($l_c$) from the ultra-thin section were quantified using ImageJ2 software, initially developed by Wayne Rasband at the National Institutes of Health

**Methods S5** Furthermore, the $A_n/C_i$ curve and chlorophyll fluorescence data at a leaf temperature of 25°C for the following species were collected: seven gymnosperms species, namely *Thuja plicata*, *Chamaecyparis obtuse*, *Juniperus oxycedrus*, *Taxus cuspidata*, *Taxus baccata*, *Sequoiadendron giganteum*, and *Picea glauca* (abbreviated as Thu, Cha, Jun, Tax_cus, Tax_bac, Seq, and Pic, respectively) used in a study by Carriquí *et al*. (2020); five ferns species, namely *Osmunda regalis*, *Marsilea quadrifolia*, *Nephrolepis exaltata* Schott, *Davallia canariensis* Sm., and *Phlebodium aureum* J.Sm (abbreviated as Osm, Mar, Nep, Dav, and Phl, respectively), and four herbs species, *Spinacia oleracea*, *Solanum lycopersicum*, *Flaveria robusta*, and *Helianthus annuus* (abbreviated as Spi, Sol_lyc, Fla, and Hel, respectively) used in a study by Nadal *et al*. (2018).

**Methods S6** For the $T_{optA}$–$T_{opt\_gm}$ correlation analysis, data on $T_{optA}$ and optimum temperature for $g_m$ ($T_{opt\_gm}$) estimated using the carbon isotope discrimination method ($g_{m\_}{}^{13}C$) in following species were collected: *Nicotiana tabacum* L. cv Petit Havana by Evans *et al*. (2013); *Oryza sativa* L. cv. Wenlu-4 and Amaroo and *Triticum aestivum* L. cv. Mace by Li *et al*. (2020); *Oryza* *sativa* L. cv. Amaroo, *Oryza sativa* meridionalis Ng, and *Oryza sativa* australiensis Domin by Scafaro *et al*. (2011); *Panicum bisulcatum* by Sonawane *et al*. (2021); and *Oryza sativa* L. cv. IR-64, *Nicotiana tabacum* cv. Petit Havana, *Gossypium hirsutum*, *Glycine max*, *Eucalyptus pauciflora*, *Lophostemon confertus*, *Quercus engelmannii*, *Triticum aestivum*, and *Arabidopsis thaliana* by von Caemmerer and Evans (2015). We also included data estimated using the chlorophyll fluorescence–gas exchange method in *Quercus canariensis* (abbreviated as Que_can) by Warren and Dreyer (2006) and *Oryza sativa* cv. Unkwang at vegetative and grain-filling stages and *Oryza sativa* cv. IR-2793 (abbreviated as Ory_Un_Veg, Ory_Un_GF, and Ory_IR, respectively) by Xue *et al*. (2016).

**Methods S7** Root mean square error (RMSE) and Nash-Sutcliffe efficiency (NSE) coefficients were used to quantify the performance of the model (Supporting Information Methods S6); the smaller the RMSE value, the better is the performance of a model. The model exhibited superior prediction performance when RMSE was lower than 25% of the mean of observations. The NSE values ranged from negative infinity to 1. An NSE value closer to 1 indicates that the model is of good quality and high reliability, whereas an NSE value closer to 0 indicates that the simulation result is close to the average level of observations and that the model is credible but with numerous simulation errors.

$$RMSE = \sqrt{\frac{1}{N}\sum_{t=1}^{N}(observed_t - simulated_t)^2} \qquad (Eqn\ S12)$$

$$NSE = 1 - \frac{\sum_{t=1}^{N}(observed_t - simulated_t)^2}{\sum_{t=1}^{N}\left(observed_t - \overline{observed}\right)^2} \qquad (Eqn\ S13)$$

where $N$ is the total number of observations; $observed_t$ and $simulated_t$ are the *t-th* observation and simulation values; $\overline{observed}$ is the mean of observations.
